# Transposable elements are associated with the variable response to influenza infection

**DOI:** 10.1101/2022.05.10.491101

**Authors:** Xun Chen, Alain Sarabia Pacis, Katherine A Aracena, Saideep Gona, Tony Kwan, Cristian Groza, Yen Lung Lin, Renata Helena Monteiro Sindeaux, Vania Yotova, Albena Pramatarova, Marie-Michelle Simon, Tomi M. Pastinen, Luis Barreiro, Guillaume Bourque

## Abstract

Influenza A virus (IAV) infections are frequent every year and result in a range of disease severity. Given that transposable elements (TEs) contribute to the activation of innate immunity, we wanted to explore their potential role in this variability. Transcriptome profiling in monocyte-derived macrophages from 39 individuals following IAV infection revealed significant inter-individual variation in viral load post-infection. Using ATAC-seq we identified a set of TE families with either enhanced or reduced accessibility upon infection. Of the enhanced families, 15 showed high variability between individuals and had distinct epigenetic profiles. Motif analysis showed an association with known immune regulators in stably enriched TE families and with other factors in variable families, including KRAB-ZNFs. We also observed a strong association between basal TE transcripts and viral load post infection. Finally, we built a predictive model suggesting that TEs, and host factors regulating TEs, contribute to the variable response to infection.

## INTRODUCTION

Influenza A virus (IAV) infection causes seasonal epidemics worldwide and results in a wide range of disease severity between individuals. The underlying reasons for this variability remain largely elusive (Clohisey and Baillie, 2019; Fukuyama and Kawaoka, 2011) but are determined by viral and host factors (Gounder and Boon, 2019). Indeed, viral determinants alone cannot account for the varied responses observed in individuals challenged by the same virus (Ciancanelli et al., 2016; Clohisey and Baillie, 2019; Gounder and Boon, 2019). The human innate immune system, which involves the modulation of several cellular pathways, is a critical component of the response to infection (Iwasaki, 2012). Upon sensing of a virus such as IAV by recognition receptors, including RIG-I and TLR3, several signal transduction pathways are triggered which further modulate various transcription factors (Bierne et al., 2012; Paschos and Allday, 2010; Xu et al., 2020). These regulators, including NF-kB/RELs, IRFs, and STATs, will engage the immune transcriptional network through the alteration of chromatin state, and in turn mediate the differential expression of hundreds of genes involved in the pro-inflammatory and antimicrobial programs to restrict virus replication and transmission (Smale, 2012; Zhang and Cao, 2021). Host factors involved in this cascade likely contribute to the variable response to IAV infection. Other factors also associated with influenza pathogenesis and that influence the response include pre-existing immunity, age, sex, obesity, and the microbiome (Gounder and Boon, 2019; Keenan and Allan, 2019). Yet, whether there exist other host factors that are important in determining the response to infection remains unknown.

Transposable elements (TEs), which occupy half of the human genome, play critical roles as cis-regulatory elements in various human biological processes (Bourque, 2009; Bourque et al., 2018; Chuong et al., 2017). Notably, a particular subclass of TEs, Endogenous Retroviruses (ERVs), are derived from ancient retrovirus, suggesting a potential association with infection and immunity (Buttler and Chuong, 2021; Kassiotis and Stoye, 2016; Srinivasachar Badarinarayan and Sauter, 2021). Confirming this, an ERV family, MER41, was found to be co-opted as cis-regulatory elements in the primate innate immune response (Bogdan et al., 2020; Chuong et al., 2016). TEs are also drastically upregulated in human immune cells upon extracellular stimuli, including viral infection (Macchietto et al., 2020; Mikhalkevich et al., 2021; Nellåker et al., 2006; Schmidt et al., 2019; Wang et al., 2020). Meanwhile, loss of SETDB1 or SUMO-modified TRIM28, which are associated with histone methylation and Kruppel-associated box domain (KRAB) zinc finger proteins (ZNFs), will lead to the significant derepression of TEs in the immune response (Cuellar et al., 2017; Schmidt et al., 2019). Together, these studies suggest that TEs play a prominent role in human innate immunity. Moreover, given that many TE families have integrated after the divergence of primates from other mammals (Benton et al., 2021) and are polymorphic in humans (Bourque et al., 2018), they could represent host factors contributing to the variable response to infection. Indeed, TE transcription is linked with aging (Bogu et al., 2019; Gorbunova et al., 2021; LaRocca et al., 2020) and microbiota (Lima-Junior et al., 2021), which are associated with the response to infection (Gounder and Boon, 2019; Keenan and Allan, 2019).

To test whether TEs and associated regulators are important host factors in the variable response to infection, we used data from a multi-omics study that profiled the transcriptome and epigenome before and after IAV infection in monocyte-derived macrophages derived from 39 individuals (Aracena et al., 2022). During the course of IAV infection, the amount of viral transcripts produced is variable and has been associated with disease severity (Clohisey and Baillie, 2019; Granados et al., 2017; de Jong et al., 2006; Li et al., 2010). Moreover, the amount of viral reads observed in the macrophages post-infection can be used as a surrogate for viral load (Thorburn et al., 2015). Indeed, in a similar experimental system this metric was shown to be stable and reproducible across individuals (O’Neill et al., 2021). Notably, by studying the infected macrophages from these 39 individuals, we observed extensive variation in the levels of viral reads and discovered a set of TEs displaying high inter-individual variability in chromatin accessibility following infection. By looking for binding motifs in these variable regions we identified novel transcription factors likely contributing to the response to infection. Lastly, using TEs and these new host factors, we were able to build models that were predictive of the response to infection as measured by the amount of viral transcripts.

## RESULTS

### Many TE families are upregulated following IAV infection but few are correlated with viral load post-infection

To characterize individual differences in the response to IAV infection, we used RNA-seq data obtained from monocyte-derived macrophages of 39 individuals before and after exposure to IAV for 24 hours (**Table S1**, see Methods and Aracena et al. 2022). As expected, we observed extensive gene expression changes upon infection (**Figure 1A**). Despite the fact that all samples engaged a strong transcriptional response to infection, we noticed extensive variation in the levels of viral reads (from 3.77% to 65.7%, **Figure 1B**), suggesting varying capacity to infection and/or to limit viral replication across individuals. Consistent with this hypothesis, viral load was inversely correlated with the expression fold change (FC) of several master regulators of the innate immune response, including transcription factors (TFs, e.g., *IRF3*, *STAT2*), adaptor molecules (e.g., *MYD88*, *TICAM1*) and interferon-inducible molecules (e.g., *IFNAR1*, *IFNAR2*) (**Figure S1A**). More globally, genes for which the transcriptional response to IAV infection was found to be correlated with viral load (*R^2^* ≥ 0.3, *p* value ≤ 0.05, **Figure S1B**), were significantly enriched for pathways involved in the viral response. Similar to protein-coding genes, TE transcription levels were also significantly changed upon infection (**Figure 1A**). We inspected TE regulation at the level of families and identified 204 upregulated and seven downregulated families (|log2FC| ≥ 1, adjusted *p* value ≤ 0.001), respectively (**Figure 1C** and **Table S2**). In line with prior studies, we observed that ERVs (also known as LTRs) were the most commonly upregulated families (179 out of 204, 85.5%) and had the strongest FC (**Figure 1C** bottom).

**Figure 1.**
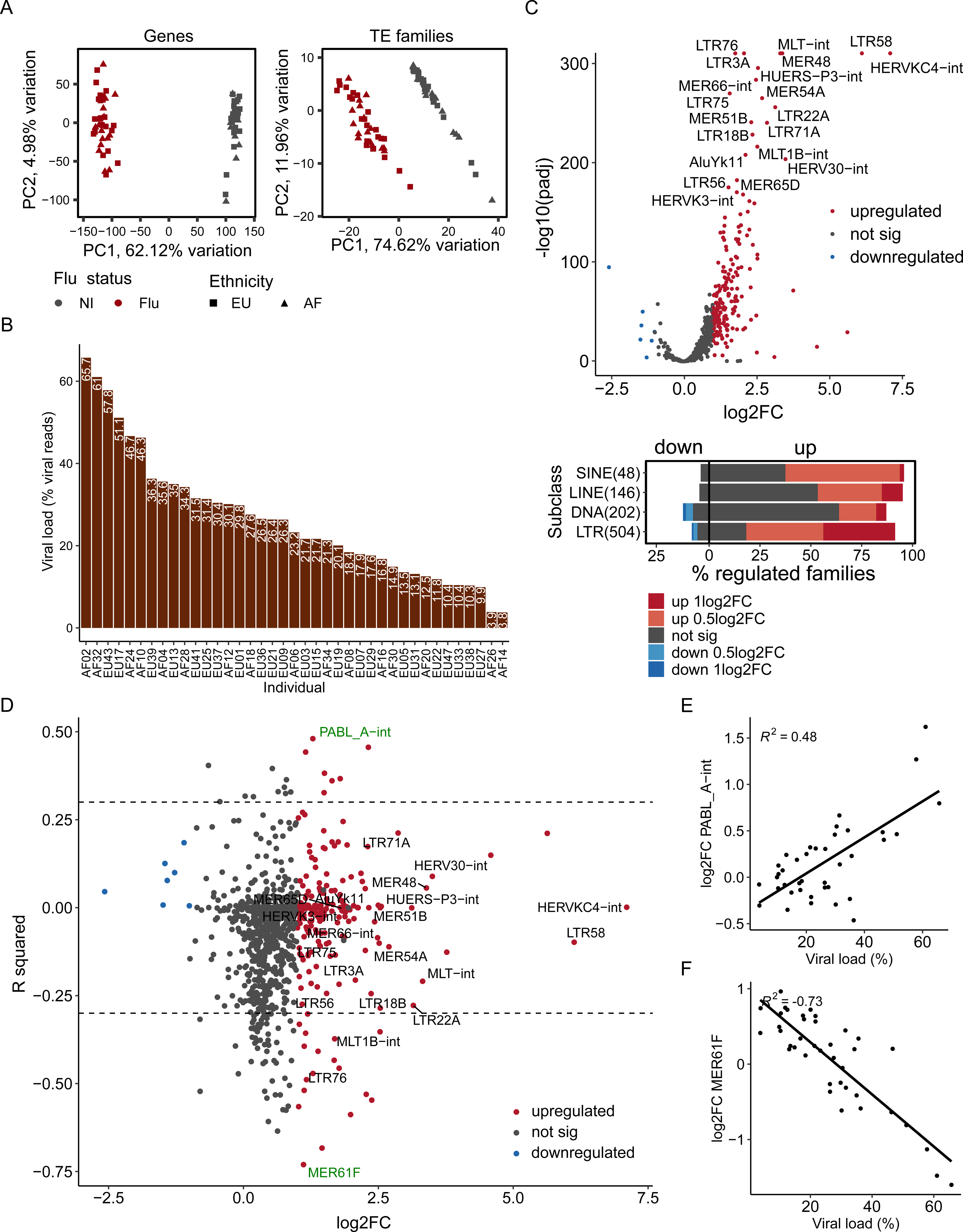
TEs are upregulated post-infection but most expression changes are not correlated to viral load. **(A)** PCA plots of genes (left) and TE families (right) expression of individuals before and after infection. Individuals with African (AF) and European (EU) ancestry are indicated. **(B)** Bar plots show viral load (% viral reads) across individuals post-infection. **(C)** TE upregulation at the family level in human macrophages in response to IAV infection. Up/down regulated families were detected as families with ≥ 1 log2 fold change (log2FC) in expression and adjusted *p* value ≤ 0.001 upon infection (top). The highest 20 upregulated families based on fold change are highlighted. The total number of examined families per TE subclass is indicated in parentheses. The vertical line separates the upregulated (left) and downregulated (right) families. **(D)** Dot plots of correlation coefficients between TE FC and viral load post-infection. X-axis represents the log2FC of each family computed by DESeq2. Y-axis represents the correlation coefficients (R squared) between expression FCs and viral load among 39 individuals. The same 20 upregulated families (Figure 1C) are highlighted here. A positively and negatively correlated family (green) is shown as examples in panel E and F respectively. **(E)** Example of positive correlation between PABL_A-int FCs and viral load. **(F)** Example of negative correlation between MER61F FCs and viral load.

Next, we looked at the correlation between TE expression FCs and viral load post-infection. Among the 902 examined families, we only identified 17 and 77 families that were positively and negatively correlated with viral load (*R*^2^ ≥ 0.3 and *p* value ≤ 0.05), respectively (**Figure 1D** and **Table S3**). For example, PABL_A-int was positively correlated with viral load (**Figure 1E**), while MER61F was negatively correlated with viral load (**Figure 1F**). Families from the LTR subclass, and ERV1 superfamily in particular, were slightly enriched for being positively correlated with viral load (**Figure S1C**). In contrast, families from the DNA subclass were more prone to negatively correlate with viral load. Taken together, we observed significant upregulation of ERVs following IAV infection but the upregulation across individuals was correlated with viral load for only a small number of repeat families.

### TEs contribute to dynamic chromatin regions in response to influenza infection

Beyond transcriptional changes, viral infection also induces significant epigenetic changes in immune cells (Zhang and Cao, 2021). We wanted to explore whether epigenetic profiles at TEs could help explain the inter-individual variability in the response to IAV infection. We used data profiling 35 of the 39 samples before and after infection using transposase-accessible chromatin using sequencing (ATAC-seq) and chromatin immunoprecipitation followed by sequencing (ChIP-seq) technologies characterizing various histone marks (**Table S1**, see Methods) (Aracena et al., 2022). Across these samples we obtained an average of 137,478 peaks for ATAC-seq, 73,190 for H3K27ac, 230,292 for H3K4me1, 33,700 H3K4me3, and 209,119 for H3K27me3 (**Figure 2A** and **Table S4**). The number of peaks across all marks was slightly higher in infected compared to non-infected samples. We observed that on average 19.5% to 47.6% of peaks were located in TEs across marks (**Figure 2B** and **Table S4**). These proportions were found to be slightly but significantly increased post-infection for H3K4me3 and H3K27me3 (student’s *t* test, *p* value ≤ 0.05). Next, to infer whether repeat regions display epigenetic variability, we measured the coefficients of variation (*cv*) in consensus peak regions (Aracena et al., 2022) and identified similar proportions of variable regions in TE and non-TE regions for most marks (0.4% to 6.4%, *cv* ≥ 0.5, **Figure 2C**, see Methods). That being said, we observed higher variability of H3K4me3 and lower variability of H3K27me3 mark in TEs compared to non-TE regions, respectively. Given that H3K4me3 is typically associated with transcription, these results are consistent with some variability of TE transcription post infection.

**Figure 2.**
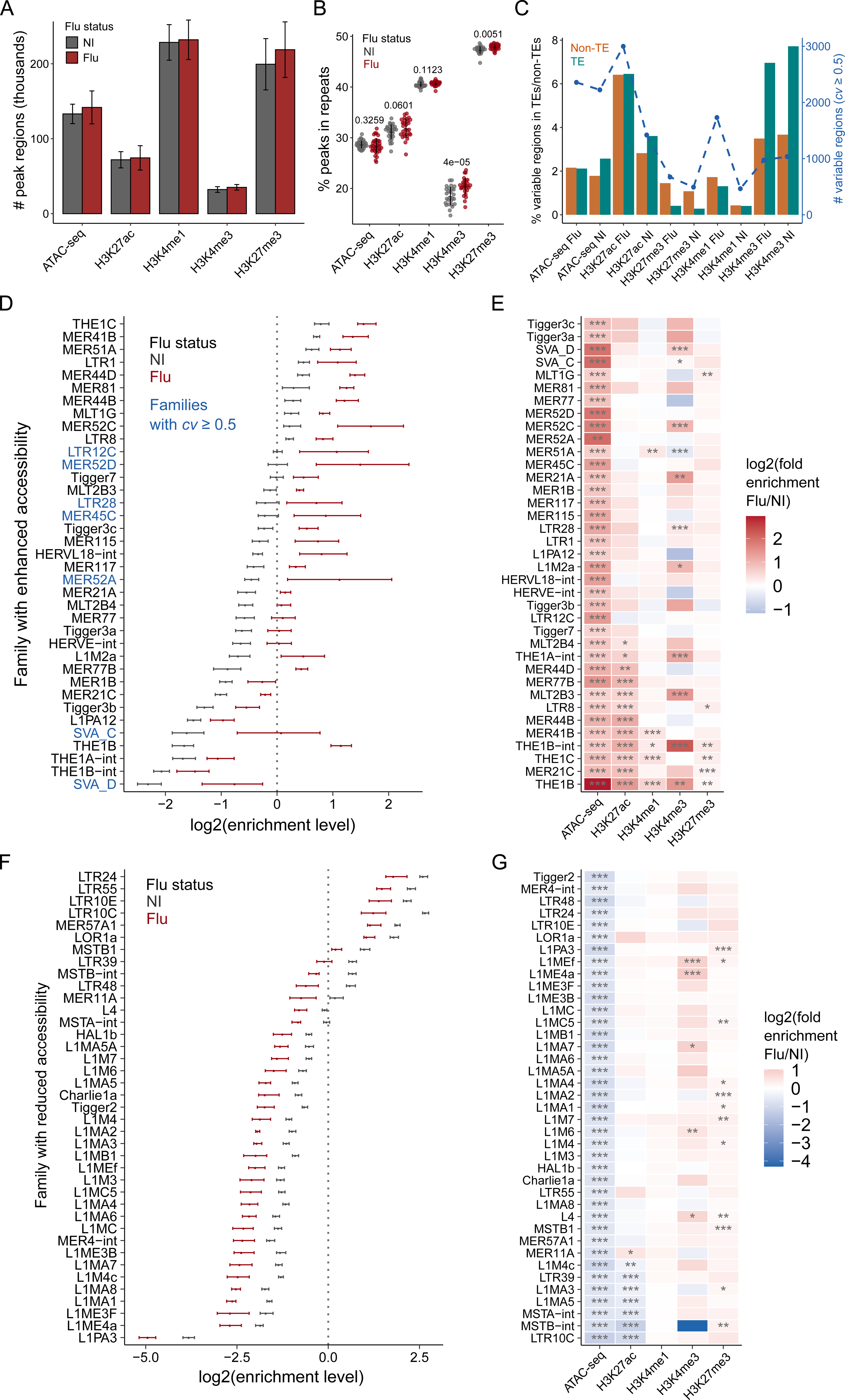
TEs contribute to dynamic chromatin regions in human macrophages in response to influenza infection. **(A)** Number of peak regions detected in infected and non-infected samples for ATAC-seq and histone marks. **(B)** Proportion of ATAC-seq and histone marks peaks that overlap repeat regions. Two-tailed paired student’s *t*-test was used to compare infected and non-infected samples for each mark. **(C)** Number and proportion of variable peak regions overlap TE and non-TE regions. Variable regions were determined with the threshold of coefficient of variation (*cv)* ≥ 0.5 (see Methods). Bars represent the proportions of peak regions that are variable while the dotted line represents the corresponding peak counts. Infected (Flu) and non-infected (NI) samples are shown separately. **(D,F)** Distribution of log2 enrichment levels of families with enhanced (D) and reduced (F) accessibility in infected and non-infected samples. Candidate families were identified using the optimized methodology as we described in **Figure S3C**. The enrichment level refers to the fold enrichment per sample relative to the corresponding random distribution (see Methods). Families with a high variability of enrichment levels between individuals (*standard deviation* divided *by the mean value, cv* ≥ 0.5) are highlighted in blue color (**Table S5**). The dotted line at “0” represents the random distribution. Standard deviations were computed in non-infected and infected samples separately. **(E,G)** Heatmap of log2 fold enrichments (Flu/NI) of families with enhanced (E) and reduced (G) accessibility for ATAC-seq and each histone mark, *i.e.*, H3K27ac, H3K4me1, H3K4me3, and H3K27me3. The fold enrichment was computed by dividing the average normalized number of peaks-associated instances in infected by non-infected samples. Two-tailed paired student’s *t*-test was used to compute the *p* values (* *p* ≤ 0.05, ** *p* ≤ 0.01, *** *p* ≤ 0.001).

To explore the TE families with accessibility changes upon IAV infection, we compared the normalized number of accessible instances per family as measured by ATAC-seq in infected versus non-infected samples (**Figure S2A**). We identified 37 families with enhanced accessibility exhibiting 1.5-fold (adjusted *p* value ≤ 0.05) or greater abundance of peaks-associated instances in infected relative to non-infected samples (**Figure S2B** and **Table S5**). For instance, we observed on average 584.2 peaks overlapping the THE1B repeat family in the flu samples, while only 79.5 were observed in the uninfected samples. The enrichment observed in these families can also be visualized relative to a random genomic background (**Figure 2D**) and include MER41B that was previously reported in K562, Hela, and CD14+ cell lines (Chuong et al., 2016). Notably, some families displayed a high degree of variation between samples post-infection (e.g. LTR12C, highlighted in blue). A similar analysis revealed that enhanced families were also frequently enriched for histone modifications, especially H3K27ac and H3K4me3 (**Figure 2E**). For instance, many H3K27ac peaks overlapped with THE1B and MER41B in infected samples (**Figure S2C**).

One of the advantages of comparing two conditions is that we could also look for TE families showing reduced accessibility upon infection. We identified 39 such families (**Figure 2F**, **Figure S2D** and **Table S5**). For instance, although on average 54.3 peaks overlapped L1M4c in non-infected samples, this number dropped to 26.0 in infected samples. Notably, 24 of the 39 (61.5%) reduced accessibility families were LINEs. This contrasts with the fact that only two out of 37 (1.7%) enhanced families were LINEs. While some families with enhanced accessibility showed high variability between individuals, families with reduced accessibility displayed a uniform profile across most individuals (**Figure 2F**). Lastly, by inspecting the enrichments of other histone modifications, we identified seven families with reduced H3K27ac (**Figure 2G** and **Table S5**). Taken together, these results highlight that a large number of epigenetically changing regions of the human genome upon IAV infection are in TEs.

### A number of TE families display high inter-individual variability upon infection

Metaplots and heatmaps of chromatin accessibility further supported the higher variability observed in some of the enhanced families post-infection. For instance, upon infection, THE1B (**Figure 3A** and **Figure S3A**) showed less variation in chromatin accessibility across individuals than LTR12C (**Figure 3B** and **Figure S3A**). To better understand why, we performed semi-supervised clustering analysis of the chromatin accessibility of the 37 enhanced families among the 35 infected samples (**Figure 3C**). This analysis revealed three groups of individuals post-infection. One outlier sample (EU37), was observed to consistently have the lowest fraction of reads in peaks (FRiP) scores among both infected and non-infected samples, suggesting a technical artifact rather than a biologically distinctive response to flu. Using this approach, a total of 15 enhanced families had the highest variability (**Figure 3C**, bottom), which we defined as “high var. families”, especially between Group 1 and Group 3 individuals. In contrast, 22 enhanced families showed consistent enrichment patterns between three individual groups, and were defined as “low var. families”. A similar analysis in the non-infected samples did not reveal any groupings, suggesting an association specific to IAV infection (**Figure S3B)**. Group 3 individuals tended to be slightly older and present higher viral loads as compared with other groups but the differences were not statistically significant (**Figure S3C-S3D**).

**Figure 3.**
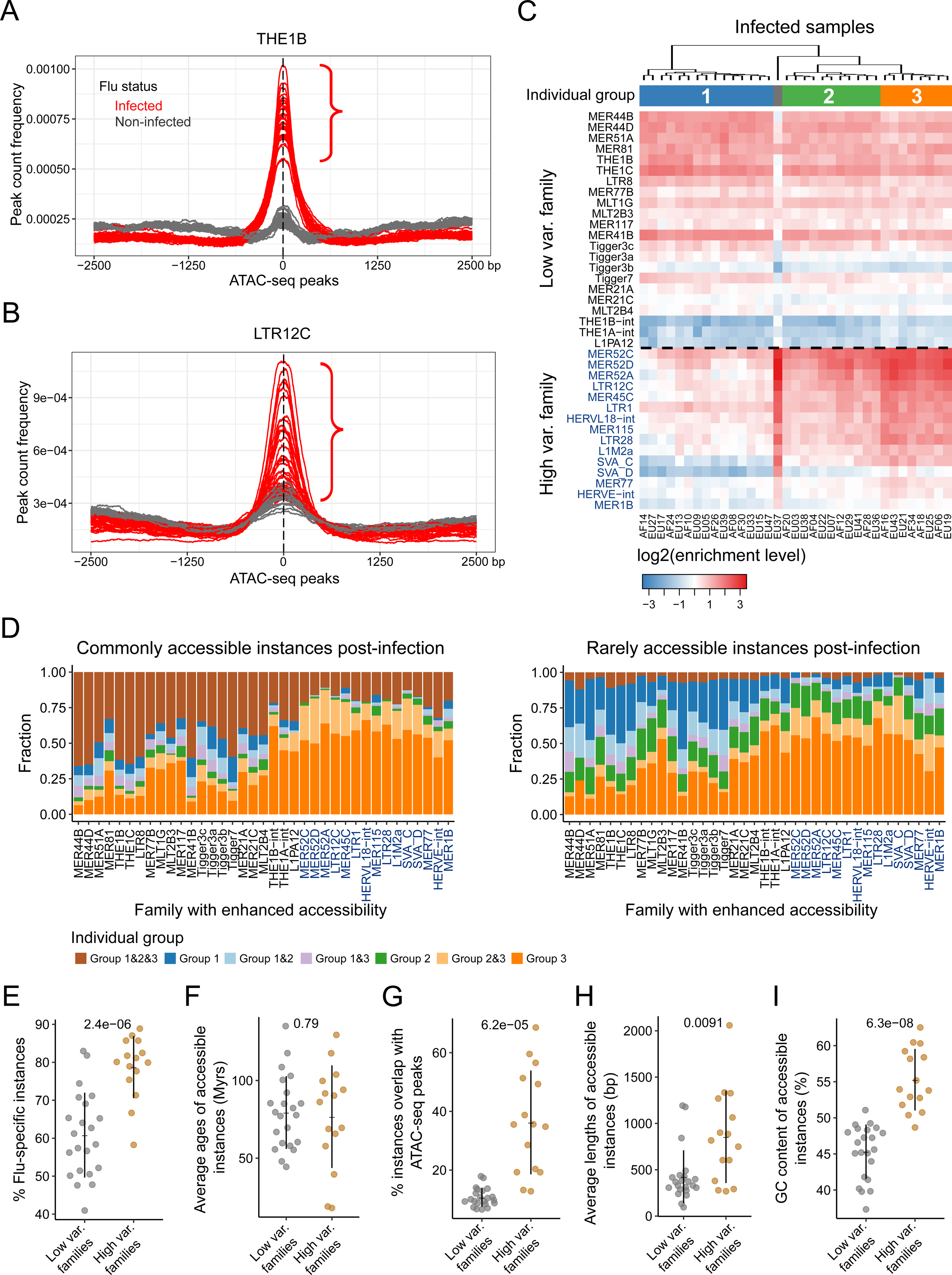
Uncovering a set of TE families that display high individual variability in chromatin accessibility post-infection. **(A-B)** Peak count frequency of ATAC-seq peaks overlapped with THE1B (A) and LTR12C (B). Red and grey lines represent the infected or non-infected samples. Compared to THE1B, LTR12C shows a higher standard deviation between infected samples. Peaks overlapping each TE instance are centered at the median position of peak summits across samples. Upstream and downstream regions (2.5 kb) are shown. **(C)** Heatmap of log2 enrichment levels of 37 families with enhanced accessibility in 35 infected samples. Semi-supervised clustering analysis was performed. Three individual groups are shown with an outlier sample. High var. families are highlighted in blue color and have higher enrichment levels in Group 3 individuals than Group 1 individuals. Enrichment level refers to the abundance of accessible instances in infected samples relative to the background. **(D)** Proportions of accessible instances per enhanced family are variable between three individual groups post infection. Commonly accessible instances represent instances that are accessible in more than 25% samples from at least one group (left); rarely accessible instances represent instances that are accessible in less than 25% samples from any groups (right). Enrichment in one individual group refers to instances that are accessible in more than 25% samples for commonly accessible instances and one or more samples for rarely accessible instances. High var. families are highlighted in blue color. **(E-I)** Comparative analysis of the proportion of flu-specific instances among all accessible instances (E), evolutionary ages (F), proportion of accessible instances among all instances (G), lengths (H) and GC contents (I) of accessible instances between high var. and low var. families. *P* values computed by two-tailed student’s *t*-test are shown above the dot plots.

Next, we asked what fraction of repeat loci from the high var. families were contributing to the variability observed between individuals. Unsupervised clustering analysis of these loci (instances) revealed that a large number displayed high variability post infection (**Figure S3E**). Among high var. families we consistently observed more commonly (≥ 25% individuals of one group) and rarely (< 25%) accessible instances that were specific to Group 3 individuals (**Figure 3D** and Methods). To further identify features that were associated with variability in accessibility in TEs, we performed a comparative analysis between high var. and low var. families. We focused on flu-specific instances (ATAC-seq peak present in ≥ 1 infected but not in non-infected samples) and found that high var. families had a significantly higher proportion as compared to low var. (student’s *t* test, *p* value = 2.4 × 10^-6^) (**Figure 3E** and **Figure S3F**). In contrast, we did not observe significant differences in the estimated evolutionary age of high var. versus low var. TE families (**Figure 3F** and **Figure S3G**). Overall, we did find that high var. families had a significantly higher proportion of instances that overlap ATAC-seq peaks, that their repeat consensus length was longer and that they had a higher GC content (**Figure 3G-3I** and **Figure S3H**). Taken together, we identified 15 TE families with increased accessibility upon infection and that have high epigenetic variability between individuals and display unique features.

### Enhanced and reduced TE families act as cis-regulatory elements in the response to influenza infection

Next, we asked if TE families with enhanced and reduced accessibility acted as cis-regulatory elements regulating nearby genes in response to IAV infection. We found that upregulated genes were more likely to be located near instances from low var. and high var. families that become accessible upon infection (flu-specific instances) (**Figure 4A**). Lower enrichments were observed for high var. compared to low var. families, indicating their weaker association to gene expression. In contrast, we observed a depletion of upregulated genes near non-infected (NI)-specific instances (accessible in ≥ 1 non-infected but not in infected samples) from TE families with reduced accessibility (**Figure 4A**). Notably, the opposite was observed for down-regulated genes (**Figure 4B**). These effects were stronger for flu-/NI-specific instances as compared to instances associated with shared peaks (**Figure S4A**). Splitting the enrichment at the TE family level, we observed consistent overrepresentation of accessible instances post-infection near upregulated genes within a 100 kb window for most enhanced families (**Figure 4C**, red color).

**Figure 4.**
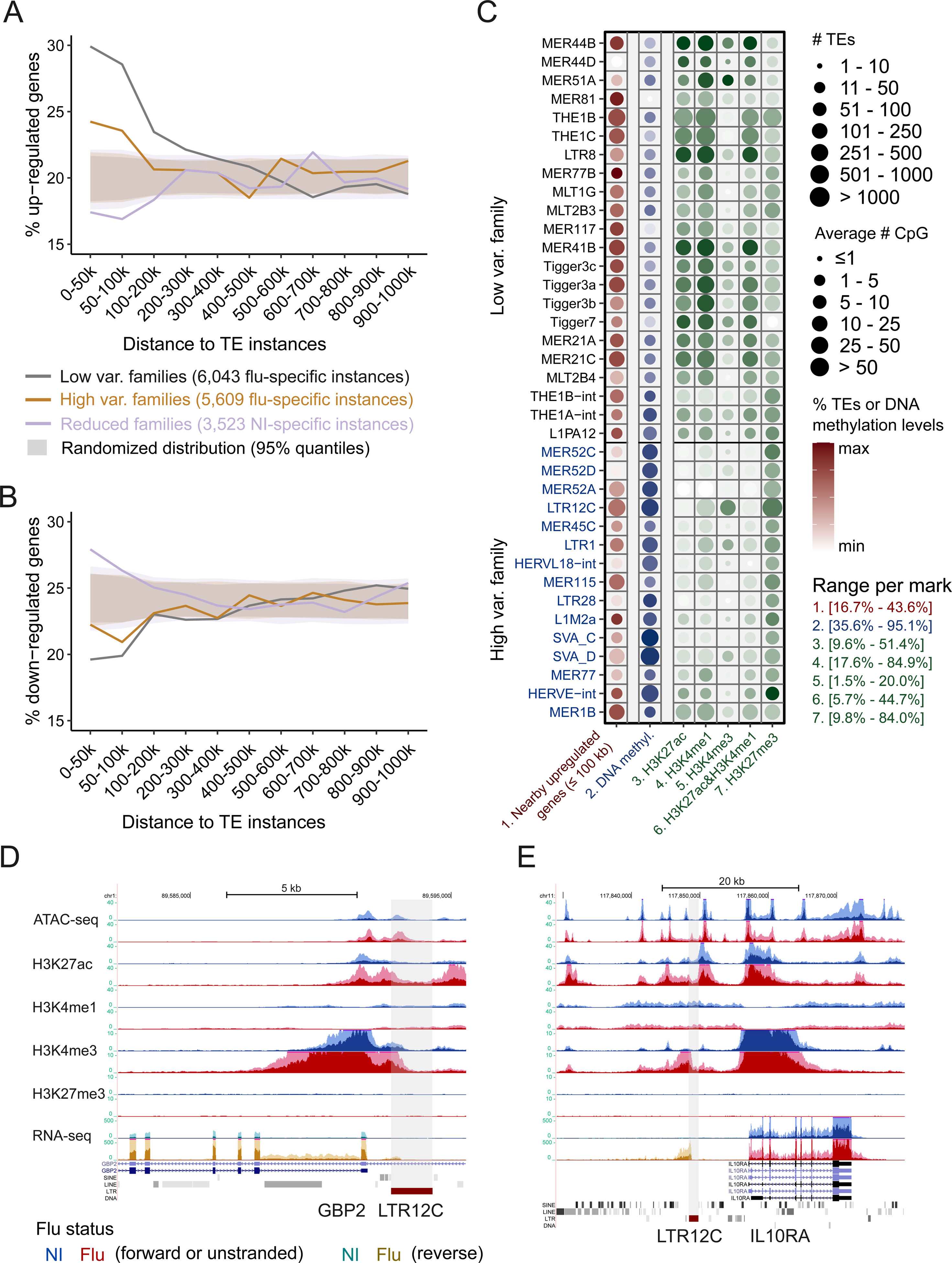
TE families with accessibility changes may play critical regulatory roles in the response to influenza infection. **(A,B)** Fractions of differentially expressed genes near accessible TEs relative to the random distributions. Proportions of up (A) and down (B) regulated genes are shown within each of the genomic intervals relative to nearby accessible TEs. Flu-specific instances from high var. and low var. families and NI-specific instances from reduced families are considered. The total number of instances are indicated in the figure legend. Expected distributions were computed by randomizing each set of accessible instances 1,000 times (shaded area, 95% confidence intervals), suggesting a statistical significance of *p* < 0.05 for values outside the distributions. The proportions of regulated genes are compared with corresponding expected distributions. **(C)** Properties of high var. and low var. families overlapped with histone marks and DNA methylation. The number and proportion of accessible instances with nearest significantly upregulated genes within 100 kb (log2FC ≥ 0.5, adjusted *p* value ≤ 0.05) are shown in red color (1st column). The number of CG sites and average DNA methylation levels are shown in blue color (2nd column). The number and proportion of accessible instances overlapped with each mark are shown in green color (3rd - 7th columns). The color ranges (proportion of accessible instances) are scaled by the minimum and maximum values for each mark. **(D)** Example genomic view of an accessible LTR12C instance potentially upregulating adjacent *GBP2* gene expression post-infection. LTR12C is highlighted as the shaded area with the increased accessibility, expression and H3K4me3 activity. The dark shaded area denotes the distribution of the average RPM values and the light shaded area denotes the standard deviation. Signals of various epigenetic marks are shown in blue color for non-infected samples and red color for infected samples. For RNA-seq, forward and reverse transcripts are shown in blue and green color separately for non-infected samples; while forward and reverse transcripts are shown in red and brown color separately for infected samples. **(E)** Example genomic view of an accessible LTR12C with the expression was upregulated and initiated at the open chromatin region post-infection. The LTR12C instance highlighted as the shaded area shows an upregulated accessibility, expression, and H3K4me3 activity. *IL10RA* gene located near the LTR12C instance is also significantly upregulated (log2FC = 1.44, adjusted *p* value = 1.60^-70^) post-infection.

Next, we investigated the properties of chromatin post infection more broadly by examining DNA methylation (**Figure 4C**, blue color) and sets of histone modifications (**Figure 4C**, green color). Instances from high var. families were highly DNA methylated (an average of 83.8%) and prone to overlap with H3K27me3 (47.3%), meanwhile they had a relatively small fraction of accessible instances overlapped with active marks (e.g. 15.1% for H3K27ac and 31.4% for H3K4me1). In contrast, low var. families were highly enriched for active histone marks (33.2% for H3K27ac and 60.7% for H3K4me1). Overall, low var. and high var. showed distinctive chromatin patterns post infection.

Finally, to further investigate which genes were potentially regulated by these TE-embedded sequences upon infection, we performed a pathway enrichment analysis using the list of nearby differentially expressed genes (≤ 100 kb). We observed an enrichment in various immune-related pathways (**Figure S4B**). For example, an LTR12C instance with enhanced chromatin accessibility accompanied by an augmentation of H3K27ac upon infection can be found in the promoter of *GBP2* (**Figure 4D**). *GBP2* gene is an interferon-induced gene and exhibits antiviral activity against IAV infection (Tretina et al., 2019). In a different LTR12C instance near the up-regulated immune-related gene *IL10RA*, transcription was initiated at the open chromatin region within the repeat itself and was flu-specific (**Figure 4E**). We also confirmed the chromatin change at the MER41 instance that was shown to be an enhancer regulating *AIM2* (**Figure S4C**) (Chuong et al., 2016). Lastly, we identified several immune-related genes that were potentially regulated by adjacent instances from enhanced families, such as the TE gene pairs of THE1C-*IFI44*, THE1C-*GBP3*, THE1B-*PSMA5*, MLT2B3-*CLEC4E*, and THE1C-*ABCG1* (**Figure S4C**). Thus, the enhanced and reduced TE families behave like cis-regulatory elements regulating nearby immune genes.

### High var. families contribute transcription factor binding sites for potentially novel host factors in the response to infection

To look for regulatory proteins associated with enhanced and reduced families, we aggregated the reads in open chromatin regions across samples to fine-map the actual peak summit on each TE instance, which was termed a “centroid”. A small fraction of instances with inaccurate or inconsistent annotations were discarded, this mostly affected TE families with reduced accessibility (**Figure S5A**). As examples, we can visualize the peak centroids identified along the consensus sequences for THE1B, a high var. family (**Figure 5A**), and LTR12C, a low var. family (**Figure 5B**). We observed a higher complexity of open chromatin regions for LTR12C compared to THE1B. Centroids were mainly detected at around 180 bp for THE1B and were scattered between 150 to 600 bp for LTR12C. Next, we defined a “TE peak region” as a location within a TE that has a peak centroid in ≥ 5 instances, starting with the region with the largest number of instances, named Region 1, and so on. For most families, more than 80% of instances were accessible in one of the top 5 TE peak regions (**Figure 5C**, inset). The location of these TE peak regions can be shown on their consensus sequence and reveals that they are quite dispersed (**Figure 5C**). Notably, compared to low var. families, high var. families had significantly more TE peak regions (student’s *t* test, *p* value = 0.022) and lower proportions of accessible instances in the top TE peak region (student’s *t* test, *p* value = 0.0037) (**Figure S5B**). This is consistent with the longer length of high var. families (**Figure 3H**).

**Figure 5.**
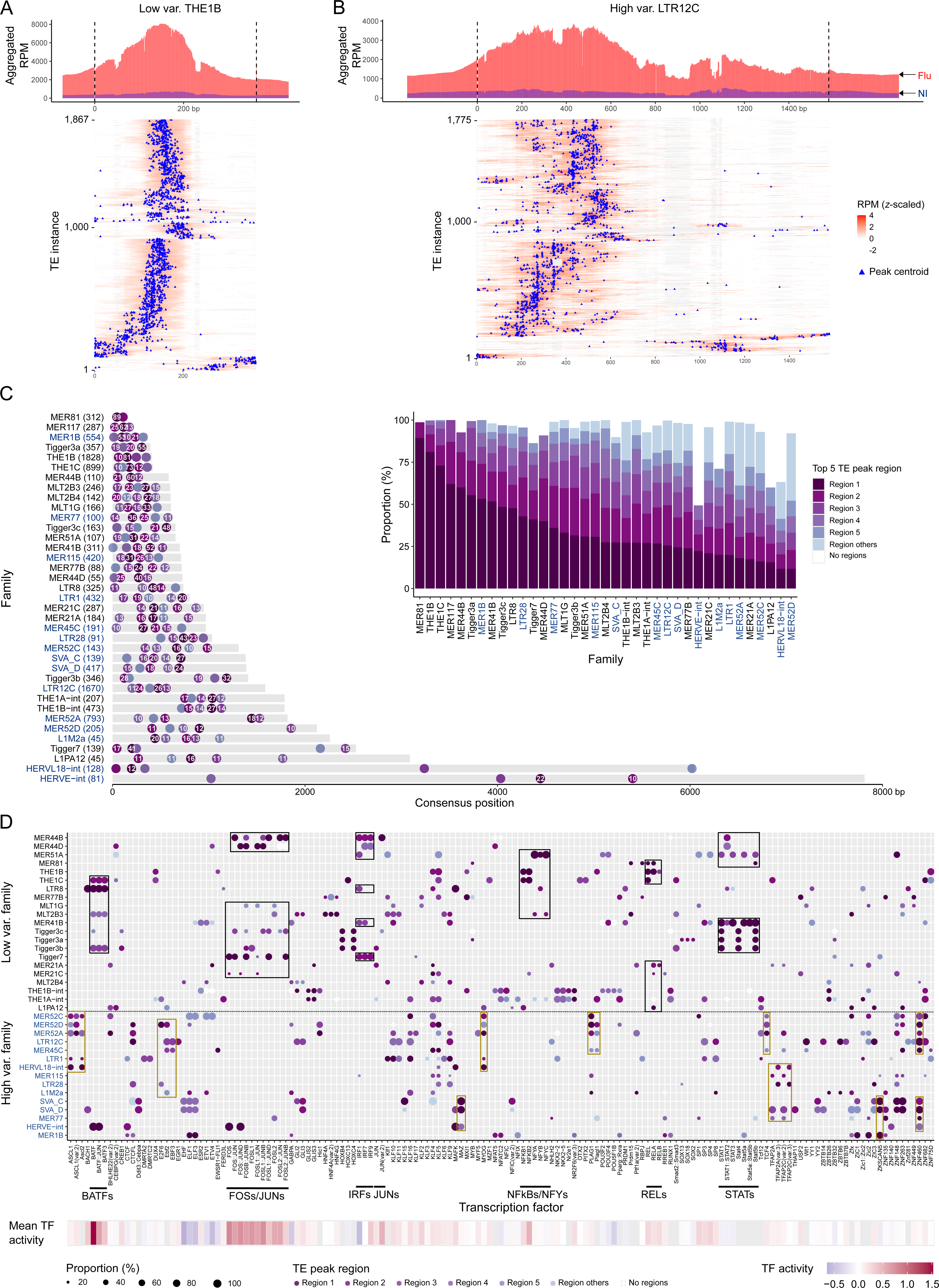
Low var. and high var. families contribute binding sites for distinct sets of potential host factors in the response to infection. **(A,B)** Distribution of chromatin accessibility along the THE1B (A) and LTR12C (B) consensus sequence. Distribution plots (up) show aggregated (summed) reads per million (RPM) values across accessible instances. Infected and non-infected samples are shown separately. Upstream and downstream regions (± 20% of the consensus sequence length) are shown. Heatmaps (bottom) show *z*-scaled RPM values per accessible instance. In the heatmap, scaled RPM values below zero are shown in white color and the deletions relative to the consensus sequence are shown in grey color. The centroid (blue triangle) refers to the peak summit per instance. The total number of instances are indicated as the y-axis. **(C)** Distribution of TE peak regions on each enhanced family. A TE peak region was previously defined as a location within a TE that has a peak centroid in ≥ 5 instances. Here, the locations and proportions (%) of the top-five TE peak regions are shown on each consensus sequence. The number in each dot refers to the proportion among accessible instances (≥ 10%) in each TE peak region. Y-axis shows the family name, consensus name, and the number of accessible instances in TE peak regions. The inset barplot shows the proportion of instances in each TE peak region. Region 1 represents the TE peak region with the highest proportion and region 2 refers to the second-highest, and so on. High var. families are in blue color. **(D)** TF binding motifs enriched in enhanced families. Same motifs enriched across TE peak regions are aggregated. TE peak regions with the most number of instances are shown as representatives. Black boxes highlighted candidate motifs recognized by known immune regulators enriched in low var. families and TF names are shown at the bottom; Brown boxes highlighted top candidate motifs recognized by potential novel host factors enriched in high var. families, including ZNF460 and ZKSCAN5. High var. families are highlighted in blue color. Mean TF activity was obtained from Aracena et al. 2022. Missing values are in grey color.

To further investigate the molecular mechanism underlying the enhanced families, we examined the TF binding motifs that were enriched in TE peak regions (**Figure 5D** and **Figure S5C**). The enrichment of binding sites for STATs and IRFs in MER41B were previously reported (Chuong et al., 2016). Here we found that the STAT related motifs mainly came from Region 1 of MER41B while IRF related motifs came from Region 3. STATs were also observed in various Tigger3 and MER44 families while IRF related motifs were also enriched in various MER44 families, LTR8 and Tigger7. Other motifs of interest observed in TE peak regions included FOS/JUN, BATFs, NFkBs/NFYs and RELs. Notably, this analysis also revealed distinct sets of binding motifs between high var. and low var. families (**Figure 5D**). Specifically, low var. families were enriched for motifs of known immune regulators; while high var. families were enriched for other motifs (e.g., ASCLs, CTCFs, EBFs, MAZ, MYOG, PLAGs, TFAP2s, ZKSCAN5, and ZNF460). We speculated that the binding of these other transcription factors may be associated with the individual epigenetic variability in high var. families post-infection. Indeed, by clustering accessible HERVE-int instances, we found that instances with peaks in Region 3 and 4, which were enriched for TFAP2 and ZNF460 motifs, were prone to be accessible in Group 3 rather than Group 1 individuals (**Figure S5D-S5E**). Supporting the potential role of KRAB-ZNFs in high var. families, we observed that binding sites for multiple ZNF TFs (Imbeault et al., 2017) were enriched in some high var. families (**Figure S5G**). ZNFs are commonly found to interact with the KAP1/TRIM28 machinery (Helleboid et al., 2019; Iyengar and Farnham, 2011). We inspected protein-protein interactions using the STRING database (Szklarczyk et al., 2019) and confirmed an association between ZKSCAN5 and TRIM28, and also between ZNF460 and TRIM28 (**Figure S5F**).

Next, we performed a similar analysis to examine the TE peak regions and corresponding motifs enriched in the 39 families with reduced accessibility (**Figure S6A-S6B**). We identified the enrichment of IRF1, MEF2A/B/C/D and SPI related motifs in these families. Notably, L1MA2, L1MA4, L1MA6, L1MA7, and L1MA8 were significantly enriched for MEF2 related motifs. MEF2 TFs are central developmental regulators (Potthoff and Olson, 2007), which are also required in the immune response that functions as an *in vivo* immune-metabolic switch (Clark et al., 2013). Lastly, by further inspecting TFs with their binding motifs that were enriched in enhanced and reduced TE families, we found that TFs bound to high var. families were mainly enriched in transcription-related pathways while TFs bound to low var. and reduced families were mainly enriched in immune-regulated pathways (**Figure S6C**). Taken together, we concluded that high var. families have a unique profile and are associated with potentially new host factors, e.g. ZNF460, which are known to be associated with the KAP1 machinery.

### TE-associated host factors can be used to predict viral load post infection

Finally, we asked whether TE and TE-associated host factors can be predictive of viral load post infection. As we previously noted, the expression changes of most TE families were not correlated with viral load (**Figure 1D**), however, we further inspected the TE expression levels in non-infected and infected samples, respectively. Unlike expression changes, we observed that the basal and post-infection expression levels of many families were correlated with viral load (**Figure 6A**, **Figure S7A** and **Table S3**). Basal expression of most TE families had comparable correlation coefficients, in contrast to post infection expression levels. Combining reads across families, we found that there was a strong inverse correlation between the total amount of basal TE transcripts and viral load post-infection (*R*^2^ = 0.45, *p* value = 2.69 × 10^-6^, **Figure 6B**). Inverse correlations were also observed for each of the four main TE subclasses (**Figure S7B**). As expected, the basal activation of the immune system (interferon signature) was also inversely correlated with viral load (**Figure 6C**, *R^2^*= 0.38, see Methods).

**Figure 6.**
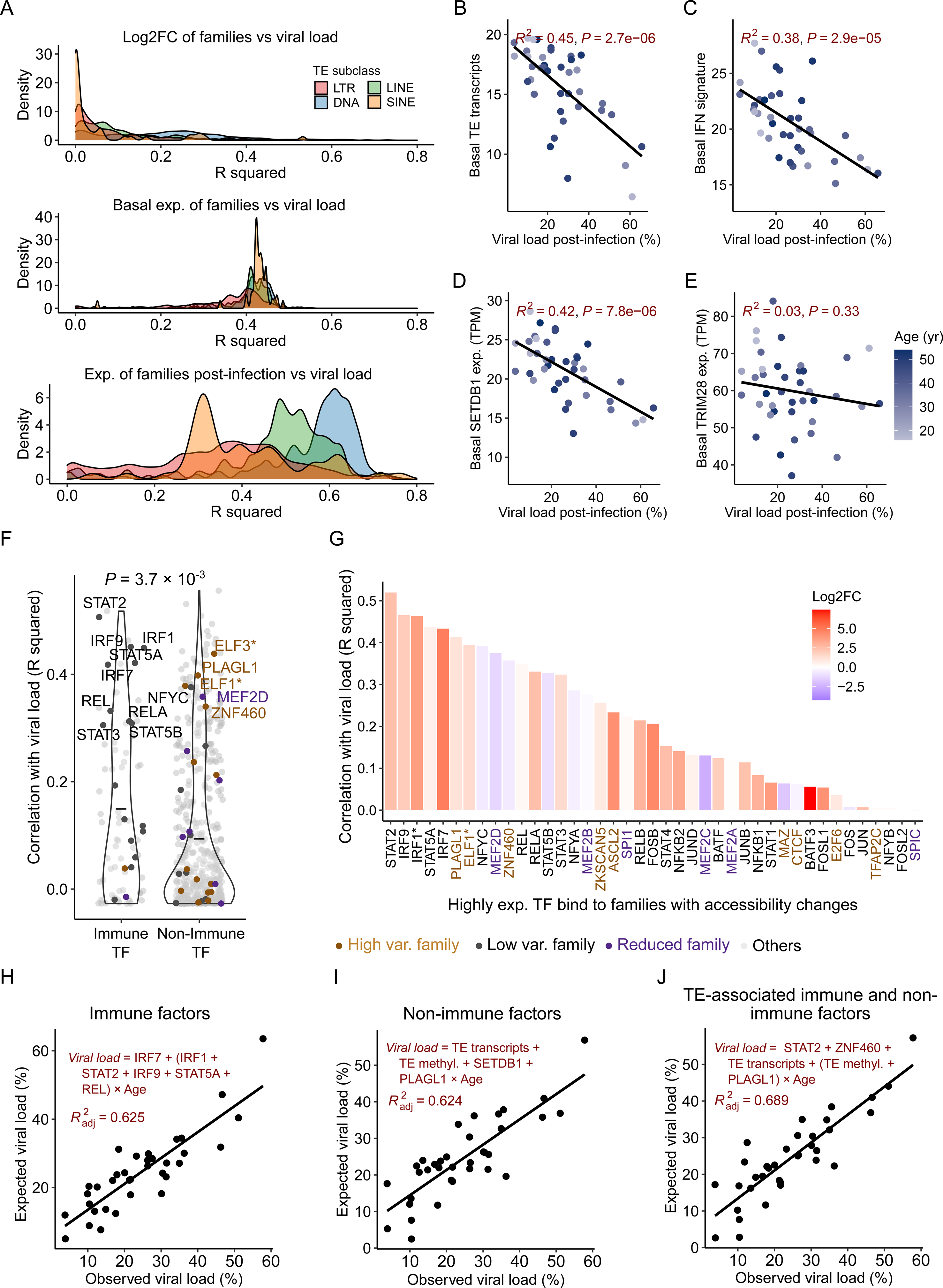
TEs and TE-associated host factors are predictive of viral load post-infection. **(A)** Distribution of correlation coefficients (R squareds) between the TE expression level (TPM) in non-infected and infected samples and TE expression fold changes with viral load post-infection. Log2FCs and TPM values were calculated as we previously described. Four TE subclasses are shown separately. Correlation directions are shown in **Figure S7A**. **(B)** Inverse correlation between the amount of basal TE transcripts and viral load. The basal TE transcript refers to the proportion of aggregated normalized read counts in TEs among the global transcripts. Black line represents the regression line. *R^2^* and *p* values computed by the linear regression model are shown. **(C)** Inverse correlation between the basal type I interferon (IFN) signature (score) and viral load. The IFN signature represents the median expression level (TPM value) of genes involved in Type I interferon signaling pathways (**Table S6**). **(D,E)** Correlations between the basal expression levels of *SETDB1* (D) and *TRIM28* (E) and viral load. It shows that *SETDB1* (*R^2^* = 0.42) rather than *TRIM28* (*R^2^* = 0.03) basal expression is associated with viral load. Basal *SETDB1* expression is also positively correlated with the basal TE transcripts and IFN signature before infection (**Figure S7G**). **(F)** Violin plot of the correlation coefficients between basal TF expression levels and viral load. Basal TPM values were used for the correlation analysis as we previously described. Immune and non-immune TFs are compared using the paired student’s *t*-test and the *p* value is also shown. Black bars represent mean values. TF genes were obtained from the JASPAR database as we previously used for the motif analysis and Immune TFs were obtained from the InnateDB database (Breuer et al., 2013). Only expressed TFs are shown. * highlights motifs that are enriched in different categories of families. **(G)** Bar plot of correlation coefficients between the basal expression of TFs bound to enhanced and reduced families. Highly expressed TFs (TPM ≥ 1) are considered and the expression fold changes upon infection are shown. TFs are ranked based on the R squared value. * highlights motifs that are enriched in different categories of families. **(H)** Multivariable regression model developed for the prediction of viral load using the expression levels of immune TFs in the basal state. The top six correlated TFs to viral load that are also associated with TEs were used. The model was generated as we described in the Methods. The formula and variables and adjusted *R^2^* are shown. **(I)** Multivariable regression model developed for the predictive of viral load using the TE-associated non-immune (novel) host factors in the basal state. Using the same approach (see Methods), a subset of features were selected among the age and six non-immune factors, including TRIM28, SETDB1, TE transcripts, TE methylation, ZNF460, and PLAGL1. **(J)** Multivariable regression model developed for the predictive of viral load using the TE-associated immune and non-immune host factors in the basal state. We included all the non-immune factors as well as STAT2 to generate the model. STAT2 was selected based on the correlation to viral load.

To explore the role of other factors known to be associated with the regulation of TEs, we inspected both *TRIM28* and *SETDB1*. We first examined the FC and observed a strong correlation to viral load post-infection for *SETDB1* but not for *TRIM28* (**Figure S7C**). Similarly, an inverse correlation was observed between *SETDB1* basal expression and viral load (*R*^2^ = 0.42, *p* value = 7.83 × 10^-6^) but not for *TRIM28* (*R*^2^ = 0.026, *p* value = 0.32) (**Figure 6D-6E**). Looking at the average DNA methylation in TEs pre-infection, we did not observe a correlation with viral load (**Figure S7D**). Age is another factor that is potentially associated with TEs, even though it was not observed to correlate with viral load (**Figure S7E**). We noted that the variability of basal TE transcription increased as the age increased (**Figure S7E**). Actually, the inverse correlation observed between basal TE transcripts and viral load became even stronger (*R*^2^ = 0.76, *p* value = 4.6 × 10^-7^) with the exclusion of individuals older than 40 years old (**Figure S7F**).

We then expanded the analysis to look at the host factors that are associated with epigenetic variability in high var. families. First, we examined the correlations between basal expression levels of all expressed TFs and viral load (**Figure 6F**). As expected, known immune-related TFs had higher correlation coefficients with viral load compared to non-immune TFs (*p* value = 3.7 × 10^-3^). Focusing on TFs associated with enhanced and reduced TE families, we found that many were strongly correlated with viral load (**Figure 6G**). From motifs found in the high var. families, we identified *PLAGL1* and *ZNF460* as the candidates with the highest correlation to viral load (**Figure S7G**, *R^2^* = 0.41 and 0.36, respectively). Notably, *PLAGL1*, which is a family member of PLAG1, also encodes a C2H2 zinc finger protein that could be repressed by SUMOylation (Dyck et al., 2004).

Lastly, we wanted to test our ability to combine all this information into predictive models to estimate the variable responses to IAV infection among the 35 individuals for which we had all the multi-omic datasets. We started with IFN related features as variables including the IFN signature and age to achieve a model explaining 36% of the variation (**Figure S7H**). Next, we included the top six immune factors bound to low var. families that were correlated with viral load as variables and used a stepwise approach to select the final set of features in a generalized linear model (see Methods). Age was also included as an interaction term variable due to its influence on multiple variables. Using this approach, we were able to build a better model (adjusted *R^2^* = 0.625) (**Figure 6H**). Afterwards, we looked at all the TE-related host factors described above in a correlation matrix chart with viral load (**Figure S7I**). Notably, when we included six non-immune factors associated with TEs and age in our model, we obtained a comparable fit with a model that includes TE transcripts and the new factor *PLAGL1* (adjusted *R^2^* = 0.624) (**Figure 6I**). Adding the top correlated immune TF, *i.e.*, *STAT2*, further increased the accuracy of the model (adjusted *R^2^*= 0.689) (**Figure 6J**). As expected, if we used age as an independent variable in these models, the predictive accuracies decreased significantly (**Figure S7J**). Altogether, we concluded that TEs and TE-related host factors can be used to predict viral load in macrophages post-infection.

## DISCUSSION

Inter-individual variability in disease is at the core of precision medicine. By examining TE transcription and epigenetic state in macrophages derived from 39 individuals, we provided new insights into the contribution of TEs to the response to IAV infection. Specifically, we discovered a set of 15 TE families with high inter-individual variability in chromatin accessibility post-infection (**Figure 3C**). Besides the distinct sequence features and chromatin states they promote, we found that high var. families mainly contribute transcription factor binding sites (TFBSs) for potentially new host factors in the response to infection (e.g., ZNF460 and ZKSCAN5); in contrast, other TE families of interest mainly contribute TFBSs for known immune regulators (**Figure 7**). Given that many of the TFBSs enriched in high var. families were associated with proteins that are known to interact with the KAP1/TRIM28 machinery, this suggests that KRAB-ZNFs may contribute to the inter-individual epigenetic variability post infection. We speculate that the enhanced accessibility in these families may be because of gradual chromatin depression led by the reduced expression of *SETDB1* or *TRIM28* upon infection.

**Figure 7.**
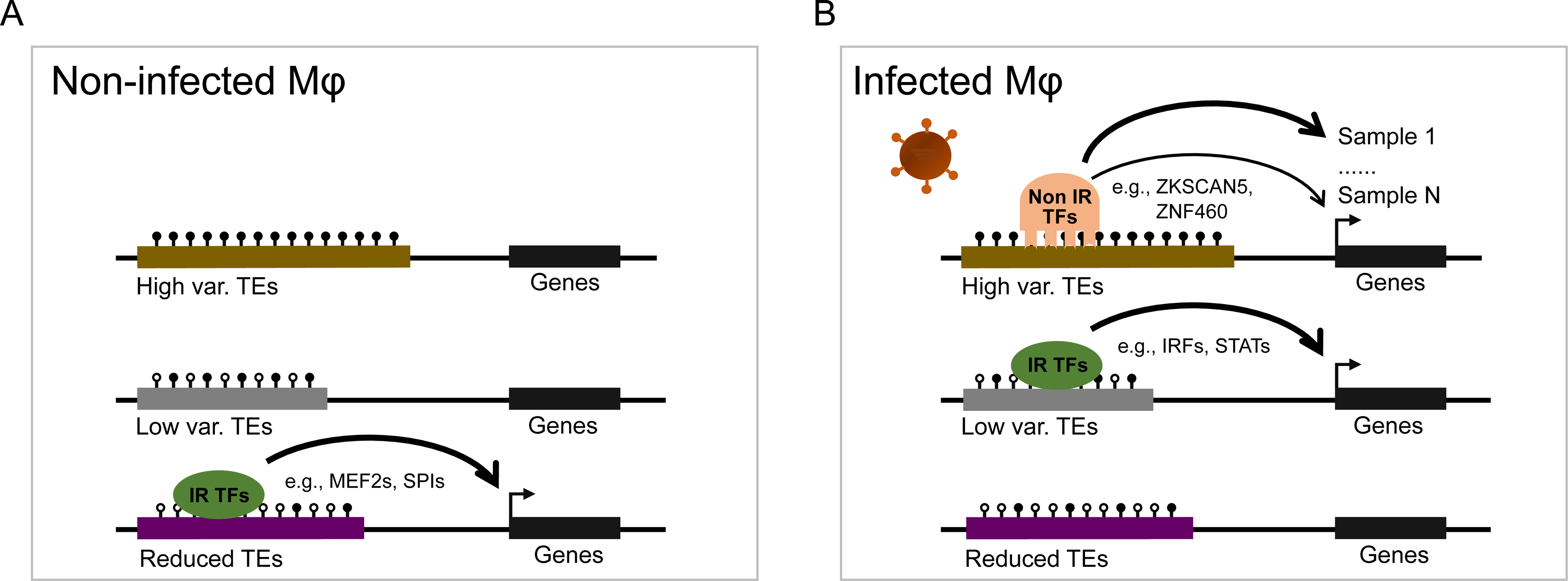
Regulatory models of TEs in response to influenza infection in human primary macrophages. **(A)** Epigenetic states of enhanced and reduced families in macrophages pre-infection. Before infection, high var. and low var. families are not accessible due to the lack of corresponding TFs binding or repression by high DNA methylation or histone methylation. In contrast, reduced families are accessible and bound by a distinct set of known immune-related (IR) TFs, including MEF2s and SPIs. High var. families are relatively longer and show a higher DNA and histone methylation level compared with other families. **(B)** Epigenetic states of enhanced and reduced families in macrophages post-infection. Chromatin accessibility of high var. and low var. families are enhanced post infection. High var. families are mainly bound by potential novel host factors (Non IR TFs), including ZFN460 and ZKSCAN5; low var. families are mainly bound by known immune-related regulators (IR TFs), including IRFs and STATs. Reduced TEs are prone to be less accessible due to the decreased expression of various TFs (e.g. MEF2s) post-infection. High var. families display a high variability in accessibility post-infection and may differentially regulate nearby genes between individuals.

In this study, multiple chromatin regions were identified for each TE family (**Figure 5C-5D**). For example, we observed the top peak region of MER44D to be significantly enriched for FOS/JUN related motifs, while another region was mainly enriched for IRF related motifs. Thus, the same TE family appears to contribute multiple binding regions recognized by different TFs, suggesting that each family may play complex regulatory roles upon infection. Additionally, by comparing the TE enrichment levels between infected and non-infected monocyte-derived macrophages following IAV infection, we were able to identify families with reduced chromatin accessibility (**Figure 2F**). These families would have been missed by previous approaches that relied on an expected distribution as control (Bogdan et al., 2020; Chuong et al., 2016; Ito et al., 2017; Sakashita et al., 2020). Moreover, although many LINE families were found to have reduced accessibility post-infection, we still observed two LINE families (L1PA12 and L1M2a) with enhanced accessibility. This may be due to the absence in these two LINE families of TFBS found enriched in their counterparts with reduced accessibility (SPIs and MEF2s).

Our data also revealed a strong inverse correlation between the basal TE transcripts and viral load post-infection. In line with the involvement of TE transcripts in the activation of innate immunity (Cuellar et al., 2017; Rookhuizen et al., 2021; Schmidt et al., 2019), we speculate that TE regulation in macrophages before infection may be involved in the activation of the innate immune response to IAV infection. To further support this claim, we combined TE basal expression levels with other factors identified in the analysis of high var. families, such as TE DNA methylation, *SETDB1* and *PLAGL1* expression levels, and were able to build a model that was predictive of the response to infection (**Figure 6H-J**). Some polymorphic TEs were also found to be eQTLs for genes upon infection, such as *TRIM25* (Groza et al., 2021), thus we speculate that polymorphic TEs may further contribute to the variable response to infection. More samples will be needed to improve and validate the predictive model we constructed using TEs and TE-associated host factors.

Altogether, our data depict major epigenetic shifts in TEs in human macrophages upon infection -- opening mostly in LTR/ERVs and closing in LINEs --, suggesting their critical role in the response to influenza infection. It is intriguing to consider that TEs might not only be an important source of regulatory innovation between species (Bogdan et al., 2020; Chuong et al., 2016) but also of regulatory variation within a population. It will be interesting to expand this analysis and study the contributions of TEs in other immune cells, e.g. CD4^+^ T cells, pneumocytes and dendritic cells (Iwasaki, 2012; Marasca et al., 2022) and to challenges with other pathogens.

## DATA AND CODE AVAILABILITY

All datasets used in this study have been deposited (Aracena et al., 2022), and are available at the European Genome-phenome Archive (EGA) as follows: RNA-seq & ATAC-seq & ChIP-seq - EGAD00001008422; and WGBS - EGAD00001008359. We also constructed a versatile browser (https://computationalgenomics.ca/tools/epivar), which allows users to explore genomic tracks for gene expression, chromatin accessibility, histone modifications, DNA methylation.

Scripts for main analyses are available at https://github.com/xunchen85/Variability_In_TEs and will be deposited at Zenodo repository once the article is accepted. Any additional information required to reanalyze the data reported in this paper is available from the lead contact upon request.

## ACKNOWLEDGEMENTS

This work was supported by a Canada Institute of Health Research (CIHR) program grant (CEE-151618) for the McGill Epigenomics Mapping Center, which is part of the Canadian Epigenetics, Environment and Health Research Consortium (CEEHRC) Network. GB is supported by a Canada Research Chair Tier 1 award, a FRQ-S, Distinguished Research Scholar award and by the World Premier International Research Center Initiative (WPI), MEXT, Japan. The Canadian Center for Computational Genomics (C3G) is supported by a Genome Canada Genome Technology Platform grant. We would like to acknowledge Calcul Québec and Compute Canada for access to computing resources. We thank Drs. Taka Inoue and Erwin Schurr for the helpful discussion and constructive comments.

## AUTHOR CONTRIBUTIONS

GB, LB and TMP planned the project and designed the experiments. RHMS, VY, AP and MMS performed all experiments in the lab. TK provided logistical support. ASP and KA performed primary data analysis and quality control. SG, CG and YLL performed some complementary data analyses. XC and GB designed all analyses presented in this study. XC performed the analyses and prepared all figures. XC and GB wrote the manuscript with the help of LB. All authors reviewed the final text.

## SUPPLEMENTAL INFORMATION

Supplementary Information includes 7 figures and 6 tables.

## METHODS

### Materials and sequencing data generation

To study the inter-individual variability in TEs following influenza A (IAV) infection, we collected primary macrophage cells from peripheral blood mononuclear cells of 39 healthy female individuals with African-American (n=19) and European-American (n=20) ancestry between 18 and 54 years old. We then infected macrophages (cultured for 6 days) with IAV for 24-hours and collected both non-infected and infected macrophages for multiple sequencing assays. The details were described here (Aracena et al., 2022). Briefly, we conducted the ATAC-seq assay to study chromatin accessibility. Using chromatin immunoprecipitation sequencing (ChIP-seq) technology, we also investigated the genome-wide profiles of H3K27ac, H3K4me1, H3K4me3, and H3K27me3 histone modifications. H3K27ac and H3K4me1 have been widely used to mark enhancers; H3K4me3 mark has been associated with promoters or active transcription; H3K27me3 mark has been associated with chromatin repression. Whole-genome bisulfite sequencing (WGBS) was further used to profile genome-wide DNA methylation. RNA sequencing (RNA-seq) was used to profile the transcriptome. All sequencing assays were performed in both infected and non-infected macrophages of each donor. Samples and generated sequencing datasets were summarized in **Table S1** (Aracena et al., 2022). Detailed methodologies to profile the genome-wide DNA methylation level and chromatin modifications were also described here (Aracena et al., 2022).

### RNA-seq read alignment

Trimmomatic (v0.36) was first used to trim adapter sequences with the parameters *PE -phred33 - quiet -validatePairs ILLUMINACLIP:$EBROOTTRIMMOMATIC/adapters/TruSeq3-PE.fa:2:30:15:2:true LEADING:3 TRAILING:30 MINLEN:50* (Bolger et al., 2014). After trimming off the adapters and low-quality nucleotides, high-quality paired-end RNA-seq reads were aligned against the human reference genome (hg19) using TopHat2 v2.1.1 (Kim et al., 2013). To optimize for the analysis of TE transcription, we kept multi-mapped reads with the recommended parameters *-x 100 --no-mixed* (Jin et al., 2015). Gene annotation file “*hg19.ensGene.gtf*” was obtained from https://hgdownload.soe.ucsc.edu/goldenPath/hg19/bigZips/genes/.

### Viral load calculation

To estimate the viral load, we re-aligned high-quality paired-end RNA-seq reads against the human reference genome (hg38) using TopHat2 with the default parameters. Paired-end unmapped reads were extracted from the unmapped BAM files and converted to FASTQ format using SAMtools (v1.10) *fastq* function (Li et al., 2009). Obtained FASTQ files were then reformatted using Fastq-pair (v0.3) tool with the parameter *-t 1000000* (Edwards and Edwards, 2019). Using TopHat2 with the same parameters, paired-end unmapped reads were aligned against the influenza A virus (H1N1) reference genome, which contains eight fragments including NC_002016.1, NC_002017.1, NC_002018.1, NC_002019.1, NC_002020.1, NC_002021.1, NC_002022.1, NC_002023.1. After that, we retrieved the number of reads mapped to influenza. Lastly, viral load was computed as the percentage of reads mapped to the influenza genome versus the total number of reads mapped to both human and influenza reference genomes.

### Gene and TE family expression measurement

TEcount implemented by TEtranscripts (v2.1.4) (Jin et al., 2015) was used to measure the gene and TE expression at the family level using RNA-seq data. Expression of each family represents the total number of reads mapped to all instances from the same family. We ran it with the use of sorted BAM file as the input and following parameters: *--sortByPos --TE hg19_rmsk_TE.gtf -- GTF hg19.ensGene.gtf --stranded reverse --mode multi*. The repeat annotation file “*hg19_rmsk_TE.gtf*” was downloaded from http://labshare.cshl.edu/shares/mhammelllab/www-data/TEtranscripts/TE_GTF/. After running, we obtained the output file for each sample which contains two columns, one column specifying the names of genes and TE families, and another column specifying corresponding read counts. The output files of all samples were combined into a count matrix for the downstream analysis.

### Gene and TE family differential expression and PCA analysis

To perform the differential expression analysis, the obtained count matrix was used as the input to DESeq2 v3.9 (Love et al., 2014). Non-infected samples were used as the control group and infected samples were used as the case group. After the removal of non-expressed TE families and genes (< 2 reads across samples), the count matrix was then standardized following QC steps of *DESeqDataSetFromMatrix*, *estimateSizeFactors*, *estimateDispersions*, and *nbinomWaldTest* included by DESeq2. Lastly, after we retrieved the output using the *results* function, we kept the significantly differentially expressed genes and TE families from DNA, LINE, SINE, LTR and SVA subclasses with the thresholds of |log2FC| ≥ 1 and adjusted *p* value ≤ 0.001.

To perform the principal component analysis (PCA), we applied a variance stabilizing transformation (vst) to the achieved normalized count matrix. We then used the PCAtools *pca* function with the parameter *removeVar = 0.1* for the PCA analysis and *biplot* function for the visualization (https://github.com/kevinblighe/PCAtools). Genes and TE families were analyzed separately.

### Gene and TE family expression levels normalization

Transcripts per kilobase million (TPM) values were calculated using the raw count matrix for genes and TE families. Specifically, we first computed the reads per kilobase (RPK) for each gene and family. For genes, we divided the read counts by the aggregated total lengths of exons per gene in kilobases; for TE families, we divided the read counts by the aggregated lengths across all instances per family. We next counted up the RPK values of both genes and TE families and divided them by 1,000,000 to obtain the TPM values.

### Correlation analysis between genes and viral load post-infection

We then examined which differentially expressed genes (DEGs) are correlated with viral load. Here, we only considered highly-expressed genes with an average of TPM values ≥ 1 in either infected or non-infected samples. The expression fold change (log2FC) of each gene was computed using the formula: *log2(TPM^Flu^+0.01) - log2(TPM^NI^+0.01).* FCs were correlated with viral load post-infection using R *lm* function. DEGs correlated with viral load (*R^2^* ≥ 0.3 and *p* value ≤ 0.05) were then submitted to the g:Profiler (https://biit.cs.ut.ee/gprofiler/gost) with the default parameters for the pathway enrichment analysis (Raudvere et al., 2019). G:SCS threshold with a minimum *p* value of 0.05 was used to determine the enriched pathways. Kyoto Encyclopedia of Genes and Genomes (KEGG) database was used to determine the enriched pathways and the top 30 terms were visualized. Key immune regulators involved in the RNA viral signaling pathway were obtained here (Xu et al., 2020). Similarly we also correlated the basal gene expression (TPM) with viral load.

### Correlation analysis between TE family and viral load post-infection

To measure the variability of TE transcription, we correlated expression fold changes of each family with viral load post-infection. Expression FC of each family per sample was computed with the same formula: *log2(TPM^Flu^+0.01) - log2(TPM^NI^+0.01)*. Similarly, R *lm* function was used for the correlation analyses. Positive and negative correlated (*R^2^* ≥ 0.3 and *p* value ≤ 0.05) families were reported.

To study the enrichment of positively or negatively associated families among each TE subclass, we performed the permutation test by comparing the actual proportion of positively/negatively correlated families among each TE subclass or superfamily relative to 10,000 randomized proportions. *P* value was calculated using the formula in R: *2 × mean (randomized_counts* ≥ *actual_count)*.

Using the same approach, we correlated the expression of TE families in infected and non-infected samples with viral load post-infection. Computed TPM values were used for the correlation analysis.

### Detection of peaks-associated TEs

After profiling the epigenetic state, we obtained ATAC-seq and Chip-seq narrow peaks in BED format. Peak regions were then converted to peak summits (median positions). To identify ATAC-seq peaks-associated instances, peak summits were intersected with the obtained repeat annotation file “hg19_rmsk_TE.gtf” using BEDtools v2.29.2 *intersect* function (Quinlan and Hall, 2010) with the parameters *-wa -u*. The same analysis was performed for other histone marks.

### Evaluation of the epigenetic variability in TEs

Unique ATAC-seq consensus peaks were obtained as we previously described (Aracena et al., 2022). To identify consensus peaks in TEs, we first converted peak regions to summits (median positions) and then intersected with the repeat annotation file aforementioned using BEDtools *intersect* function with the parameters *-wa -wb*. After that, read counts were normalized to RPM value for the downstream comparative analysis across samples. Specifically, the read count was first divided by the total number of reads and then multiplied 1,000,000. The coefficient of variation (*cv*) of each peak region was computed using the formula: *cv = absolute(sd/mean)*. Infected and non-infected samples were analyzed separately. Consensus peak regions with a minimum RPM value of “1” were kept. Variable regions were defined as the peak regions with *cv* values ≥ 0.5, referring to regions with the standard deviation that is half of the mean. Proportions of variable regions in TEs and non-TEs were analyzed separately. Same analysis was performed for other histone marks.

### Detection of TE families with chromatin state changes

We next aimed to identify families with enhanced accessibility upon infection. Firstly, we normalized the number of peaks-associated instances per family. Briefly, we divided the number of peaks-associated instances by the total number of peaks per sample, and then multiplied the average number of peaks across samples. Infected and non-infected samples were normalized, separately. Secondly, to identify families with enhanced accessibility during infection, we kept families with significantly more peaks-associated instances (≥ 1.5-fold, adjusted *p* value ≤ 0.05) in infected than non-infected samples. Two-tailed paired student’s *t*-test was used for the comparison and the resulting *p* value was adjusted for multiple testing with the Benjamini-Hochberg using the R *p.adjust* function. Lastly, we kept family candidates from DNA, LINE, SINE, LTR, and SVA subclasses with a minimum of 20 peaks-associated instances on average among either infected or non-infected samples.

Similarly, to identify families with reduced accessibility, we kept families with significantly more peaks-associated instances (≥ 1.5-fold, adjusted *p* value ≤ 0.05) in non-infected than infected samples. Same analysis was applied to each histone mark to identify families with dynamic regulatory (e.g., enhancer or promoter) potentials upon infection.

We also computed the enrichment level of each family by comparing the actual number of peaks-associated instances with its expected distribution (Bogdan et al., 2020). Specifically, we first annotated peaks-associated instances using BEDtools *intersect* function with the parameters *-wa -u* based on the annotation files (*i.e.*, desert, distal, proximal, 5′ untranslated region (5’UTR), promoter, transcription start site (TSS), exon, and intron regions) obtained from https://github.com/lubogdan/ImmuneTE. We then shuffled the true peaks while keeping the distribution relative to each region using BEDtools *shuffle* function with the parameters *-incl* or *-excl*, for 1000 times. The randomized peaks were intersected with the repeat annotation file to achieve the number of expected peaks-associated instances per family. Lastly, we computed the enrichment level of each family as the actual number of peaks-associated instances relative to the average number of the expected values.

### TE clustering analysis

To identify families with high variability, we performed the semi-supervised clustering analysis of enhanced families in 35 infected samples. Here, to rule out the impacts of different genomic distribution between TE families, we used the enrichment level relative to the expected distribution rather than the actual number of instances for the clustering analysis. Briefly, the enrichment levels of enhanced families were gathered into a data matrix followed by the log2 conversion. R *heatmap.2* function was used to perform the unsupervised clustering analysis with the default parameters. Based on the obtained enrichment pattern among samples, we re-ordered the families. Families with higher enrichment levels in Group 3 individuals than Group 1 individuals were distinguished. Non-infected samples were analyzed separately.

We then want to understand whether individual instances from high var. families display a high variability in infected samples. Peaks-associated instances from high var. families were collected. Instances with open chromatin were recorded as “1”; instances with closed chromatin were recorded as “0”. We then performed the clustering analysis using R *hclust* function with the default parameters.

### Detection of TE instances from enhanced families with variable accessibility

For each accessible instance, we first computed the percentage of samples from each group that were accessible post-infection. Next, we defined commonly accessible instances as the instances that were accessible in 25% or more samples from one individual group; we also defined rarely accessible instances as the instances that were accessible in less than 25% samples from any groups. An instance that was accessible in more than 25% samples for commonly accessible instances and one or more samples for rarely accessible instances was considered as enriched in one individual group. Lastly, we computed the proportion of instances that were prone to be accessible in each group.

### TE age estimation

The evolutionary age of each instance was estimated using our previous approach (Bogdan et al., 2020; Bourque et al., 2008). In brief, the sequence divergence of each instance relative to the corresponding consensus sequence was obtained from the “.align” file generated by RepeatMasker (https://www.repeatmasker.org/). Hg19 “.align” file was obtained from the UCSC database (https://hgdownload.soe.ucsc.edu/goldenPath/hg19/bigZips/). The divergence rate of each instance was divided by the substitution rate for the human genome (2.2×10^-9^) to compute the age per instance (Lander et al., 2001). The average ages across all instances was referred to the age of each TE family.

### Detection of peak centroids on accessible instances

We next want to fine-map the peak centroid on each accessible instance. Read depths were extracted from the aligned BAM file using BEDtools *genomecov* function with the parameter *-d* and then divided by 1,000,000 to compute the RPM values. We then aggregated (summed) RPM values of each nucleotide across accessible instances. Infected and non-infected samples were analyzed separately. The nucleotide with the highest RPM value was recorded as the peak centroid of each instance. Peak centroids in infected samples were used for families with enhanced accessibility; peak centroids in non-infected samples were used for families with reduced accessibility.

### Sequence alignment of instances against consensus sequences

We next wanted to map accessible instances to corresponding consensus sequences. The aforementioned RepeatMasker “.align” file was used to retrieve the consensus positions at single-nucleotide resolution. Instances with consistent start and end positions with the “.out” file were kept for downstream analyses. The inconsistency was potentially due to the defective annotation methodologies for the nested instances, extremely short instances, etc. It was a fact that instances of one TE family may be aligned to different consensus sequences. Thus, we wanted to focus on instances aligned to the most representative consensus sequence for each family. In the end, we pinpointed the peak centroid to the consensus sequence.

We plotted the aggregated RPM values relative to the consensus sequence using R. We also clustered accessible instances using the RPM values relative to the consensus sequence. Specifically, after *z*-transformation, scaled RPM values ≤ 0 and consensus regions with deletions were recoded as “0”. R function *heatmap.2* with the default parameter was used for the unsupervised clustering analysis. Heatmap was plotted using *ggplot2* in R.

### Detection of TE peak regions

We next wanted to identify “TE peak regions”, which referred to the consensus regions that become accessible on multiple instances. We first excluded instances that were only accessible in the outlier sample and then used the sliding window approach to identify TE peak regions. To iterate over the entire consensus sequence, the window size was set at 100 bp with a step size of one base pair. In each step, we counted the total number of peak centroids within each 100 bp window. The 100 bp-window containing the most peak centroids was identified as a TE peak region (≥ 5 peak centroids). After the exclusion of previously counted peak centroids, the analysis was repeated till all candidate TE peak regions were identified. The proportion of instances in each TE peak region was computed. TE peak regions were identified using peak centroids in infected samples for enhanced families and non-infected samples for reduced families.

### Transcription factor binding motifs analysis within TE peak regions

Firstly, we extracted 100 bp sequence centered at the centroid of each TE instance using BEDtools *getfasta* function with the *-s* parameter and then used the MEME *fimo* function to search the extracted sequences for known motifs from the latest 8th release of JASPAR motif database (http://jaspar.genereg.net/download/CORE/JASPAR2020_CORE_vertebrates_non-redundant_pfms_meme.txt) (Bailey et al., 2009; Fornes et al., 2020). Instances uniquely accessible in the outlier sample were excluded. Secondly, instances were categorized into each TE peak region, e.g., TE peak region with the most instances was named as “Region 1” and so on. TE peak regions with less than five instances were excluded. Instances not in TE peak regions were grouped as “No regions”. Thirdly, we computed the proportion of instances (100 bp centered at the centroid) containing each motif for each TE peak region. The top 5 most abundant motifs in each TE peak region were kept as candidates. To obtain enriched motifs per family, we kept motif candidates appearing in more than 20% instances in each TE peak region and more than 50% instances per family. Lastly, the same motifs detected in multiple TE peak regions were aggregated (summed) to recalculate the proportion; motifs enriched in a total of ≥ 50 instances across families were kept as top candidates. After the analysis, enriched motifs were compared between different TE peak regions and families.

### Protein-protein interaction

To identify the protein association networks of ZNF TFs (ZNF460 and ZKSCAN5) that were associated with high var. families, we submitted them to the STRING database (https://version-11-0.string-db.org) with the default parameters.

### TE regulation of neighboring genes

To explore whether TEs regulate neighboring genes, we examined differentially expressed genes (DEGs) nearby flu-specific instances from enhanced families and nearby NI-specific instances from reduced families. After the differential expression analysis, we retrieved corresponding gene names and coordinates through the command line and parameters: *mysql --user=genome -N--host=genome-mysql.cse.ucsc.edu -A -D hg19 -e “select ensGene.name, name2, chrom, strand, txStart, txEnd, value from ensGene, ensemblToGeneName where ensGene.name = ensemblToGeneName.name”*. To compute the distance between genes and TEs, the first nucleotide (5’ end) (TSS) was used to represent each gene and the median position was used to represent each TE instance. Highly expressed genes (average TPM values ≥ 1 in either infected or non-infected samples) were used for the analysis. BEDtools *window* function was used to obtain human genes centered at each accessible instance within an 1-Mb window. We then computed the proportion of significantly upregulated and downregulated genes among inspected genes, respectively, within each interval of 0-50 kb, 50-100 kb, 100-200 kb, 200-300 kb and so on. Each gene was counted once within each interval.

We also compared the proportions of significantly up/down regulated genes with the expected distribution to compute the statistical significance. Accessible instances were randomly shuffled for high var., low var. families, and reduced families for 1000 times separately. After the detection of genes near accessible instances, the proportions of significantly up/down regulated genes were computed as the expected values. The binomial distribution of the proportions of up/down regulated genes within each genomic interval was plotted with the 95% confidence interval, suggesting a statistical significance of *p* < 0.05 for any observed values outside the distribution. We then compared the proportions of significantly up/down regulated genes near accessible instances from high var. families, low var. families, and families with reduced accessibility.

We also compared the proportion of up/down regulated genes between flu-specific, NI-specific instances and instances overlapped with shared peaks (instances that were accessible in both infected and non-infected samples).

### Profile of DNA methylation and various histone marks of accessible instances

Focusing on enhanced families, we calculated the number and proportion of accessible instances overlapped with each mark post-infection. Specifically, we used BEDtools *intersect* function to identify accessible instances overlapped with each histone mark in infected samples. The median position of each peak was used for the analysis. We further identified instances overlapped with both H3K27ac and H3K4me1 marks in infected samples, suggesting the active or strong enhancer potential. We also computed the number and proportion of nearby DEGs within 100 kb (log2FC ≥ 0.5, adjusted *p* value ≤ 0.05). Additionally, we computed the average DNA methylation level of each instance and then we used the mean value across instances to represent the DNA methylation level of the family. DNA methylation level was calculated as the number of methylated cytosines divided by the sum of methylated and unmethylated cytosines at each locus.

### Pathway enrichment analysis of genes potentially regulated by TEs

The list of significantly up/down regulated genes near each accessible instance was obtained using BEDtools2 *window* function with the parameters *-l 100000 -r 100000*. The transcription start site was used to represent each gene. We focused on the significantly upregulated genes near accessible instances for high var. and low var. families, and significantly downregulated genes near accessible instances for reduced families. The obtained gene lists were submitted to the g:profiler tool with the same settings for the pathway enrichment analysis. We visualized the enriched pathways using *ggplot2* in R.

### Calculation of the amount of global TE transcripts

The amount of global TE transcripts was computed as the proportion of aggregated (summed) read counts normalized by DEseq2 in TEs among the total RNA-seq read counts in both TEs and genes. The linear regression model was used to evaluate the correlation between the basal TE transcripts and viral load post-infection. R *lm* function was used for the analysis and the corresponding *p* value and *R^2^* were reported. Using the same approach, we further analyzed each of the four main TE subclasses, *i.e.*, DNA, LINE, SINE and LTR.

### Calculation of the average DNA methylation levels in TEs

We computed the average DNA methylation levels among examined CpG sites across all annotated TE regions (TE methylation) in non-infected samples. TE families from the four main subclasses were considered.

### Construction of predictive models for viral load post-infection

Multiple regression analysis was used to build the predictive models. Viral load post-infection was used as the outcome of the models. The baseline of IFN signature (score) was computed as the median TPM value amongst 39 expressed genes from type I IFN signaling pathways (**Table S6**). We first included the baseline of IFN signature and age as predictive variables. We then chose the top six correlated immune TFs of which basal expression levels are also associated with TEs as variables, including *STAT2*, *IRF1*, *IRF7*, *IRF9*, *STAT5A*, and *REL*. We also picked non-immune factors that were associated with TEs as predictive variables, including age, the basal amount of TE transcripts, the average DNA methylation levels in TEs (TE methylation), and the basal expression levels (TPM) of *TRIM28*, *SETDB1*, *PLAGL1*, and *ZNF460*. R *glm* function with the parameter *family = gaussian()* was first used to include all variables in the generalized linear model. R *stepAIC* function was then used to choose a subset of main features for the final model. R *summary* function was used to report the *R^2^*, adjusted *R^2^* and *p* value. Lastly, we used the R *predict* function with the parameter *type = “response”* for the expected viral load with each predictive model.

## REFERENCES

1. Aracena, K.A., Lin, Y.-L., Luo, K., Pacis, A., Gona, S., Mu, Z., Yotova, V., Sindeaux, R., Pramatarova, A., Simon, M.-M., et al. (2022). Epigenetic variation impacts ancestry-associated differences in the transcriptional response to influenza infection. Submitted.

2. Bailey, T.L., Boden, M., Buske, F.A., Frith, M., Grant, C.E., Clementi, L., Ren, J., Li, W.W., and Noble, W.S. (2009). MEME Suite: tools for motif discovery and searching. Nucleic Acids Res. 37, W202–W208. https://doi.org/10.1093/nar/gkp335.

3. Benton, M.L., Abraham, A., LaBella, A.L., Abbot, P., Rokas, A., and Capra, J.A. (2021). The influence of evolutionary history on human health and disease. Nat. Rev. Genet. 22, 269–283. https://doi.org/10.1038/s41576-020-00305-9.

4. Bierne, H., Hamon, M., and Cossart, P. (2012). Epigenetics and Bacterial Infections. Cold Spring Harb. Perspect. Med. 2, a010272. https://doi.org/10.1101/cshperspect.a010272.

5. Bogdan, L., Barreiro, L., and Bourque, G. (2020). Transposable elements have contributed human regulatory regions that are activated upon bacterial infection. Philos. Trans. R. Soc. B Biol. Sci. 375, 20190332. https://doi.org/10.1098/rstb.2019.0332.

6. Bogu, G.K., Reverter, F., Marti-Renom, M.A., Snyder, M.P., and Guigó, R. (2019). Atlas of transcriptionally active transposable elements in human adult tissues. BioRxiv 714212. https://doi.org/10.1101/714212.

7. Bolger, A.M., Lohse, M., and Usadel, B. (2014). Trimmomatic: a flexible trimmer for Illumina sequence data. Bioinformatics 30, 2114–2120. https://doi.org/10.1093/bioinformatics/btu170.

8. Bourque, G. (2009). Transposable elements in gene regulation and in the evolution of vertebrate genomes. Curr. Opin. Genet. Dev. 19, 607–612. https://doi.org/10.1016/j.gde.2009.10.013.

9. Bourque, G., Leong, B., Vega, V.B., Chen, X., Lee, Y.L., Srinivasan, K.G., Chew, J.-L., Ruan, Y., Wei, C.-L., Ng, H.H., et al. (2008). Evolution of the mammalian transcription factor binding repertoire via transposable elements. Genome Res. 18, 1752–1762. https://doi.org/10.1101/gr.080663.108.

10. Bourque, G., Burns, K.H., Gehring, M., Gorbunova, V., Seluanov, A., Hammell, M., Imbeault, M., Izsvák, Z., Levin, H.L., Macfarlan, T.S., et al. (2018). Ten things you should know about transposable elements. Genome Biol. 19, 199. https://doi.org/10.1186/s13059-018-1577-z.

11. Breuer, K., Foroushani, A.K., Laird, M.R., Chen, C., Sribnaia, A., Lo, R., Winsor, G.L., Hancock, R.E.W., Brinkman, F.S.L., and Lynn, D.J. (2013). InnateDB: systems biology of innate immunity and beyond—recent updates and continuing curation. Nucleic Acids Res. 41, D1228–D1233. https://doi.org/10.1093/nar/gks1147.

12. Buttler, C.A., and Chuong, E.B. (2021). Emerging roles for endogenous retroviruses in immune epigenetic regulation*. Immunol. Rev. 305. https://doi.org/10.1111/imr.13042.

13. Chuong, E.B., Elde, N.C., and Feschotte, C. (2016). Regulatory evolution of innate immunity through co-option of endogenous retroviruses. Science 351, 1083–1087. https://doi.org/10.1126/science.aad5497.

14. Chuong, E.B., Elde, N.C., and Feschotte, C. (2017). Regulatory activities of transposable elements: from conflicts to benefits. Nat. Rev. Genet. 18, 71–86. https://doi.org/10.1038/nrg.2016.139.

15. Ciancanelli, M.J., Abel, L., Zhang, S.-Y., and Casanova, J.-L. (2016). Host genetics of severe influenza: from mouse Mx1 to human IRF7. Curr. Opin. Immunol. 38, 109–120. https://doi.org/10.1016/j.coi.2015.12.002.

16. Clark, R.I., Tan, S.W.S., Péan, C.B., Roostalu, U., Vivancos, V., Bronda, K., Pilátová, M., Fu, J., Walker, D.W., Berdeaux, R., et al. (2013). MEF2 Is an In Vivo Immune-Metabolic Switch. Cell 155, 435–447. https://doi.org/10.1016/j.cell.2013.09.007.

17. Clohisey, S., and Baillie, J.K. (2019). Host susceptibility to severe influenza A virus infection. Crit. Care 23, 303. https://doi.org/10.1186/s13054-019-2566-7.

18. Cuellar, T.L., Herzner, A.-M., Zhang, X., Goyal, Y., Watanabe, C., Friedman, B.A., Janakiraman, V., Durinck, S., Stinson, J., Arnott, D., et al. (2017). ---Silencing of retrotransposons by SETDB1 inhibits the interferon response in acute myeloid leukemia--. J. Cell Biol. 216, 3535–3549. https://doi.org/10.1083/jcb.201612160.

19. Dyck, F.V., Delvaux, E.L.D., Ven, W.J.M.V. de, and Chavez, M.V. (2004). Repression of the Transactivating Capacity of the Oncoprotein PLAG1 by SUMOylation *. J. Biol. Chem. 279, 36121–36131. https://doi.org/10.1074/jbc.M401753200.

20. Edwards, J.A., and Edwards, R.A. (2019). Fastq-pair: efficient synchronization of paired-end fastq files. BioRxiv 552885. https://doi.org/10.1101/552885.

21. Fornes, O., Castro-Mondragon, J.A., Khan, A., van der Lee, R., Zhang, X., Richmond, P.A., Modi, B.P., Correard, S., Gheorghe, M., Baranašić, D., et al. (2020). JASPAR 2020: update of the open-access database of transcription factor binding profiles. Nucleic Acids Res. 48, D87–D92. https://doi.org/10.1093/nar/gkz1001.

22. Fukuyama, S., and Kawaoka, Y. (2011). The pathogenesis of influenza virus infections: the contributions of virus and host factors. Curr. Opin. Immunol. 23, 481–486. https://doi.org/10.1016/j.coi.2011.07.016.

23. Gorbunova, V., Seluanov, A., Mita, P., McKerrow, W., Fenyö, D., Boeke, J.D., Linker, S.B., Gage, F.H., Kreiling, J.A., Petrashen, A.P., et al. (2021). The role of retrotransposable elements in ageing and age-associated diseases. Nature 596, 43–53. https://doi.org/10.1038/s41586-021-03542-y.

24. Gounder, A.P., and Boon, A.C.M. (2019). Influenza Pathogenesis: The Effect of Host Factors on Severity of Disease. J. Immunol. 202, 341–350. https://doi.org/10.4049/jimmunol.1801010.

25. Granados, A., Peci, A., McGeer, A., and Gubbay, J.B. (2017). Influenza and rhinovirus viral load and disease severity in upper respiratory tract infections. J. Clin. Virol. 86, 14–19. https://doi.org/10.1016/j.jcv.2016.11.008.

26. Groza, C., Chen, X., Pacis, A., Simon, M.-M., Pramatarova, A., Aracena, K.A., Pastinen, T., Barreiro, L.B., and Bourque, G. (2021). Genome graphs detect human polymorphisms in active epigenomic states during influenza infection. 2021.09.29.462206. https://doi.org/10.1101/2021.09.29.462206.

27. Helleboid, P.-Y., Heusel, M., Duc, J., Piot, C., Thorball, C.W., Coluccio, A., Pontis, J., Imbeault, M., Turelli, P., Aebersold, R., et al. (2019). The interactome of KRAB zinc finger proteins reveals the evolutionary history of their functional diversification. EMBO J. 38, e101220. https://doi.org/10.15252/embj.2018101220.

28. Imbeault, M., Helleboid, P.-Y., and Trono, D. (2017). KRAB zinc-finger proteins contribute to the evolution of gene regulatory networks. Nature 543, 550–554. https://doi.org/10.1038/nature21683.

29. Ito, J., Sugimoto, R., Nakaoka, H., Yamada, S., Kimura, T., Hayano, T., and Inoue, I. (2017). Systematic identification and characterization of regulatory elements derived from human endogenous retroviruses. PLOS Genet. 13, e1006883. https://doi.org/10.1371/journal.pgen.1006883.

30. Iwasaki, A. (2012). A Virological View of Innate Immune Recognition. Annu. Rev. Microbiol. 66, 177–196. https://doi.org/10.1146/annurev-micro-092611-150203.

31. Iyengar, S., and Farnham, P.J. (2011). KAP1 Protein: An Enigmatic Master Regulator of the Genome. J. Biol. Chem. 286, 26267–26276. https://doi.org/10.1074/jbc.R111.252569.

32. Jin, Y., Tam, O.H., Paniagua, E., and Hammell, M. (2015). TEtranscripts: a package for including transposable elements in differential expression analysis of RNA-seq datasets. Bioinformatics 31, 3593–3599. https://doi.org/10.1093/bioinformatics/btv422.

33. de Jong, M.D., Simmons, C.P., Thanh, T.T., Hien, V.M., Smith, G.J.D., Chau, T.N.B., Hoang, D.M., Chau, N.V.V., Khanh, T.H., Dong, V.C., et al. (2006). Fatal outcome of human influenza A (H5N1) is associated with high viral load and hypercytokinemia. Nat. Med. 12, 1203–1207. https://doi.org/10.1038/nm1477.

34. Kassiotis, G., and Stoye, J.P. (2016). Immune responses to endogenous retroelements: taking the bad with the good. Nat. Rev. Immunol. 16, 207–219. https://doi.org/10.1038/nri.2016.27.

35. Keenan, C.R., and Allan, R.S. (2019). Epigenomic drivers of immune dysfunction in aging. Aging Cell 18, e12878. https://doi.org/10.1111/acel.12878.

36. Kim, D., Pertea, G., Trapnell, C., Pimentel, H., Kelley, R., and Salzberg, S.L. (2013). TopHat2: accurate alignment of transcriptomes in the presence of insertions, deletions and gene fusions. Genome Biol. 14, R36. https://doi.org/10.1186/gb-2013-14-4-r36.

37. Lander, E.S., Linton, L.M., Birren, B., Nusbaum, C., Zody, M.C., Baldwin, J., Devon, K., Dewar, K., Doyle, M., FitzHugh, W., et al. (2001). Initial sequencing and analysis of the human genome. Nature 409, 860–921. https://doi.org/10.1038/35057062.

38. LaRocca, T.J., Cavalier, A.N., and Wahl, D. (2020). Repetitive elements as a transcriptomic marker of aging: Evidence in multiple datasets and models. Aging Cell 19, e13167. https://doi.org/10.1111/acel.13167.

39. Li, C.-C., Wang, L., Eng, H.-L., You, H.-L., Chang, L.-S., Tang, K.-S., Lin, Y.-J., Kuo, H.-C., Lee, I.-K., Liu, J.-W., et al. (2010). Correlation of Pandemic (H1N1) 2009 Viral Load with Disease Severity and Prolonged Viral Shedding in Children - Volume 16, Number 8—August 2010 - Emerging Infectious Diseases journal - CDC. https://doi.org/10.3201/eid1608.091918.

40. Li, H., Handsaker, B., Wysoker, A., Fennell, T., Ruan, J., Homer, N., Marth, G., Abecasis, G., Durbin, R., and 1000 Genome Project Data Processing Subgroup (2009). The Sequence Alignment/Map format and SAMtools. Bioinformatics 25, 2078–2079. https://doi.org/10.1093/bioinformatics/btp352.

41. Lima-Junior, D.S., Krishnamurthy, S.R., Bouladoux, N., Collins, N., Han, S.-J., Chen, E.Y., Constantinides, M.G., Link, V.M., Lim, A.I., Enamorado, M., et al. (2021). Endogenous retroviruses promote homeostatic and inflammatory responses to the microbiota. Cell 0. https://doi.org/10.1016/j.cell.2021.05.020.

42. Love, M.I., Huber, W., and Anders, S. (2014). Moderated estimation of fold change and dispersion for RNA-seq data with DESeq2. Genome Biol. 15, 550. https://doi.org/10.1186/s13059-014-0550-8.

43. Macchietto, M.G., Langlois, R.A., and Shen, S.S. (2020). Virus-induced transposable element expression up-regulation in human and mouse host cells. Life Sci. Alliance 3. https://doi.org/10.26508/lsa.201900536.

44. Marasca, F., Sinha, S., Vadalà, R., Polimeni, B., Ranzani, V., Paraboschi, E.M., Burattin, F.V., Ghilotti, M., Crosti, M., Negri, M.L., et al. (2022). LINE1 are spliced in non-canonical transcript variants to regulate T cell quiescence and exhaustion. Nat. Genet. 54, 180–193. https://doi.org/10.1038/s41588-021-00989-7.

45. Mikhalkevich, N., O’Carroll, I.P., Tkavc, R., Lund, K., Sukumar, G., Dalgard, C.L., Johnson, K.R., Li, W., Wang, T., Nath, A., et al. (2021). Response of human macrophages to gamma radiation is mediated via expression of endogenous retroviruses. PLOS Pathog. 17, e1009305. https://doi.org/10.1371/journal.ppat.1009305.

46. Nellåker, C., Yao, Y., Jones-Brando, L., Mallet, F., Yolken, R.H., and Karlsson, H. (2006). Transactivation of elements in the human endogenous retrovirus W family by viral infection. Retrovirology 3, 44. https://doi.org/10.1186/1742-4690-3-44.

47. O’Neill, M.B., Quach, H., Pothlichet, J., Aquino, Y., Bisiaux, A., Zidane, N., Deschamps, M., Libri, V., Hasan, M., Zhang, S.-Y., et al. (2021). Single-Cell and Bulk RNA-Sequencing Reveal Differences in Monocyte Susceptibility to Influenza A Virus Infection Between Africans and Europeans. Front. Immunol. 12.

48. Paschos, K., and Allday, M.J. (2010). Epigenetic reprogramming of host genes in viral and microbial pathogenesis. Trends Microbiol. 18, 439–447. https://doi.org/10.1016/j.tim.2010.07.003.

49. Potthoff, M.J., and Olson, E.N. (2007). MEF2: a central regulator of diverse developmental programs. Development 134, 4131–4140. https://doi.org/10.1242/dev.008367.

50. Quinlan, A.R., and Hall, I.M. (2010). BEDTools: a flexible suite of utilities for comparing genomic features. Bioinformatics 26, 841–842. https://doi.org/10.1093/bioinformatics/btq033.

51. Raudvere, U., Kolberg, L., Kuzmin, I., Arak, T., Adler, P., Peterson, H., and Vilo, J. (2019). g:Profiler: a web server for functional enrichment analysis and conversions of gene lists (2019 update). Nucleic Acids Res. 47, W191–W198. https://doi.org/10.1093/nar/gkz369.

52. Rookhuizen, D.C., Bonte, P.-E., Ye, M., Hoyler, T., Gentili, M., Burgdorf, N., Durand, S., Aprahamian, F., Kroemer, G., Manel, N., et al. (2021). Induction of transposable element expression is central to innate sensing.

53. Sakashita, A., Maezawa, S., Takahashi, K., Alavattam, K.G., Yukawa, M., Hu, Y.-C., Kojima, S., Parrish, N.F., Barski, A., Pavlicev, M., et al. (2020). Endogenous retroviruses drive species-specific germline transcriptomes in mammals. Nat. Struct. Mol. Biol. 27, 967–977. https://doi.org/10.1038/s41594-020-0487-4.

54. Schmidt, N., Domingues, P., Golebiowski, F., Patzina, C., Tatham, M.H., Hay, R.T., and Hale, B.G. (2019). An influenza virus-triggered SUMO switch orchestrates co-opted endogenous retroviruses to stimulate host antiviral immunity. Proc. Natl. Acad. Sci. 116, 17399–17408. https://doi.org/10.1073/pnas.1907031116.

55. Smale, S.T. (2012). Transcriptional regulation in the innate immune system. Curr. Opin. Immunol. 24, 51–57. https://doi.org/10.1016/j.coi.2011.12.008.

56. Srinivasachar Badarinarayan, S., and Sauter, D. (2021). Switching Sides: How Endogenous Retroviruses Protect Us from Viral Infections. J. Virol. 95, e02299–20. https://doi.org/10.1128/JVI.02299-20.

57. Szklarczyk, D., Gable, A.L., Lyon, D., Junge, A., Wyder, S., Huerta-Cepas, J., Simonovic, M., Doncheva, N.T., Morris, J.H., Bork, P., et al. (2019). STRING v11: protein-protein association networks with increased coverage, supporting functional discovery in genome-wide experimental datasets. Nucleic Acids Res. 47, D607–D613. https://doi.org/10.1093/nar/gky1131.

58. Thorburn, F., Bennett, S., Modha, S., Murdoch, D., Gunson, R., and Murcia, P.R. (2015). The use of next generation sequencing in the diagnosis and typing of respiratory infections. J. Clin. Virol. 69, 96–100. https://doi.org/10.1016/j.jcv.2015.06.082.

59. Tretina, K., Park, E.-S., Maminska, A., and MacMicking, J.D. (2019). Interferon-induced guanylate-binding proteins: Guardians of host defense in health and disease. J. Exp. Med. 216, 482–500. https://doi.org/10.1084/jem.20182031.

60. Wang, M., Qiu, Y., Liu, H., Liang, B., Fan, B., Zhou, X., and Liu, D. (2020). Transcription profile of human endogenous retroviruses in response to dengue virus serotype 2 infection. Virology 544, 21–30. https://doi.org/10.1016/j.virol.2020.01.014.

61. Xu, Q., Tang, Y., and Huang, G. (2020). Innate immune responses in RNA viral infection. Front. Med. https://doi.org/10.1007/s11684-020-0776-7.

62. Zhang, Q., and Cao, X. (2021). Epigenetic Remodeling in Innate Immunity and Inflammation. Annu. Rev. Immunol. 39, 279–311. https://doi.org/10.1146/annurev-immunol-093019-123619.

