## Supplemental Figures and Table legends for "Transposable elements are associated with the variable response to influenza infection"

#### SUPPLEMENTARY FIGURES

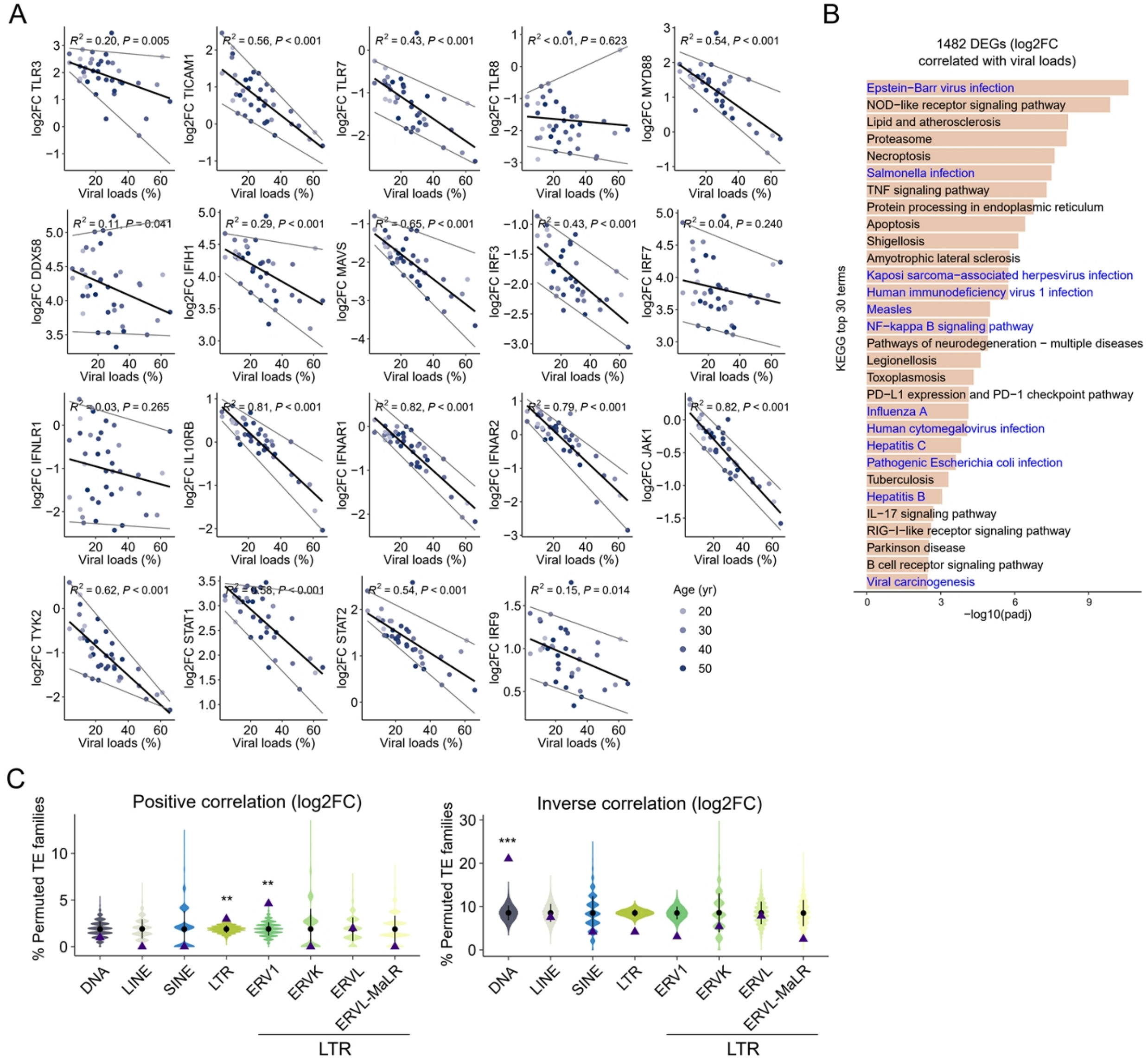


**Figure S1. Immune genes and few TE families are prone to correlate with viral load post-infection**

**(A)** Expression fold changes (log2FCs) of most key immune regulators are inversely correlated with viral load post-infection. Many genes are shown to strongly correlate with viral load, including membrane-bound receptor genes sensing infection-induced interferons, *i.e.*, *IL10RB*, *IFNAR1*, *IFNAR2*, and *JAK1*. Linear regression model was used for the correlation analysis.

Black line represents the regression line and grey lines represent the 5% and 95% quantiles. **(B)** Expression fold change amongst differentially expressed genes (DEGs) correlated with viral load are enriched in multiple virus infection pathways. 1,482 DEGs with log2FCs correlated with viral load ($R^2 \geq 0.3$, $p$ value $\leq 0.05$) were identified and used for downstream pathway enrichment analysis. **(C)** Enrichment of the proportion for log2FC amongst TE families positively and inversely correlated with viral load post-infection. 17 positively and 77 inversely correlated TE families are analyzed. Purple triangle represents the actual proportion of correlated families among subclass or superfamily. Error bar represents the mean values and standard deviations of 10,000 randomized proportions. One-tailed student's *t*-test was used to compare the actual proportions with randomized proportions ($^{*}$ $p \leq 0.05$, $^{**}$ $p \leq 0.01$, $^{***}$ $p \leq 0.001$).

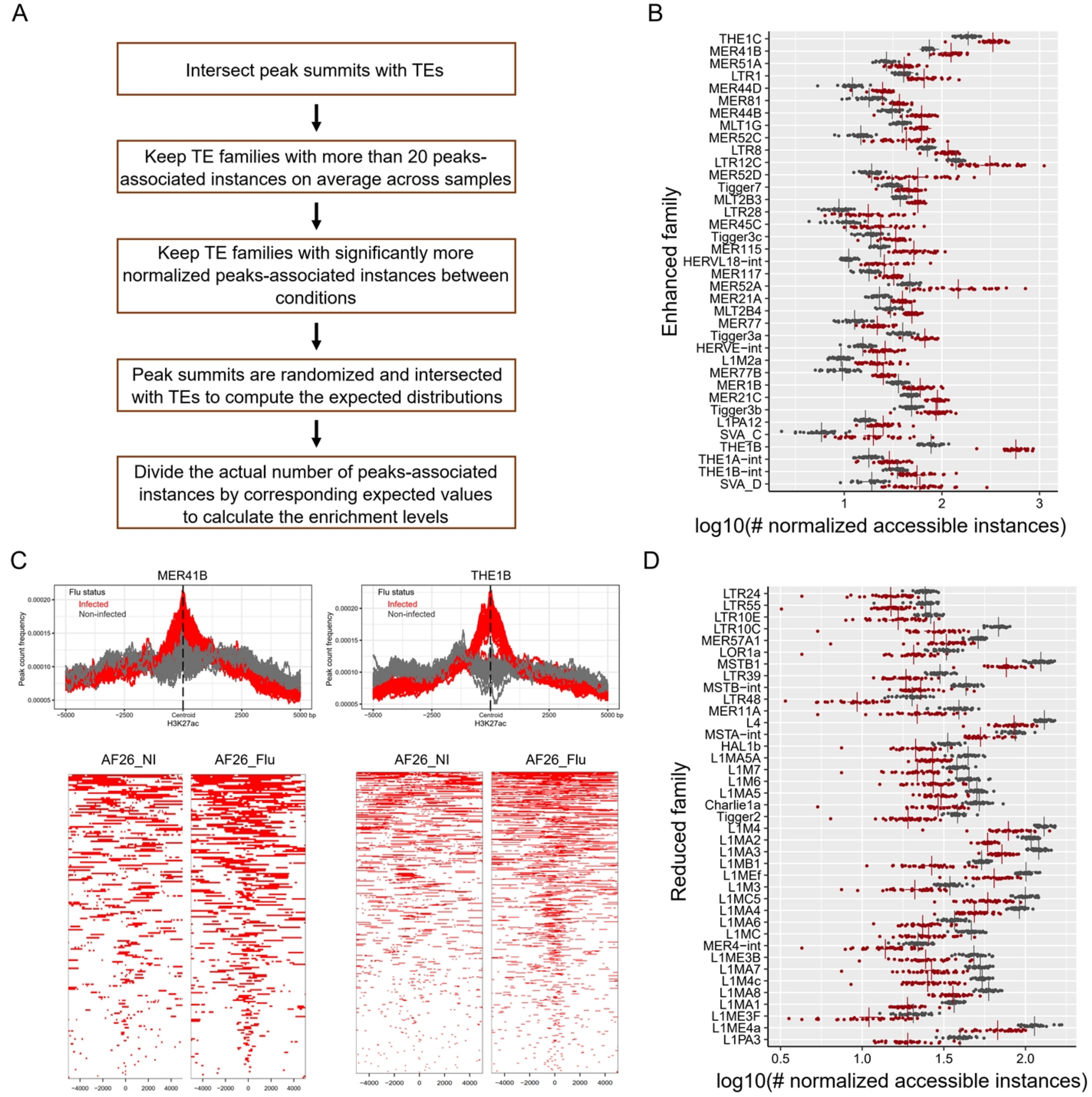


**Figure S2. Detection of TE families with enhanced and reduced accessibility in response to IAV infection**

**(A)** Optimized TE enrichment analysis pipeline. Details were described in Methods. **(B)** Distributions of the normalized number of peaks-associated instances of TE families with enhanced accessibility. Each point represents one sample. Grey color represents the non-infected sample and red color represents the infected sample. “+” indicates the mean value across non-infected (grey) and infected (red) samples. The Number of accessible instances were normalized by the average number of peaks across infected and non-infected samples separately. **(C)** Average profiles (up) and heatmaps (bottom) of H3K27ac peaks at MER41B and THE1B (± 5 kb). H3K27ac peaks are centered at the median positions of peak summits across infected samples. Peak regions are shown as heatmaps at the bottom. AF26 non-infected and infected samples are shown as examples. Peak regions centered at each family are shown as red bars. **(D)** Distributions of the normalized number of peaks-associated instances of TE families with reduced accessibility. Each point represents one non-infected (grey) and infected (red) sample. “+” indicates the mean value across non-infected (grey) and infected (red) samples. The number of accessible instances were normalized by the average number of peaks across infected and non-infected samples separately.

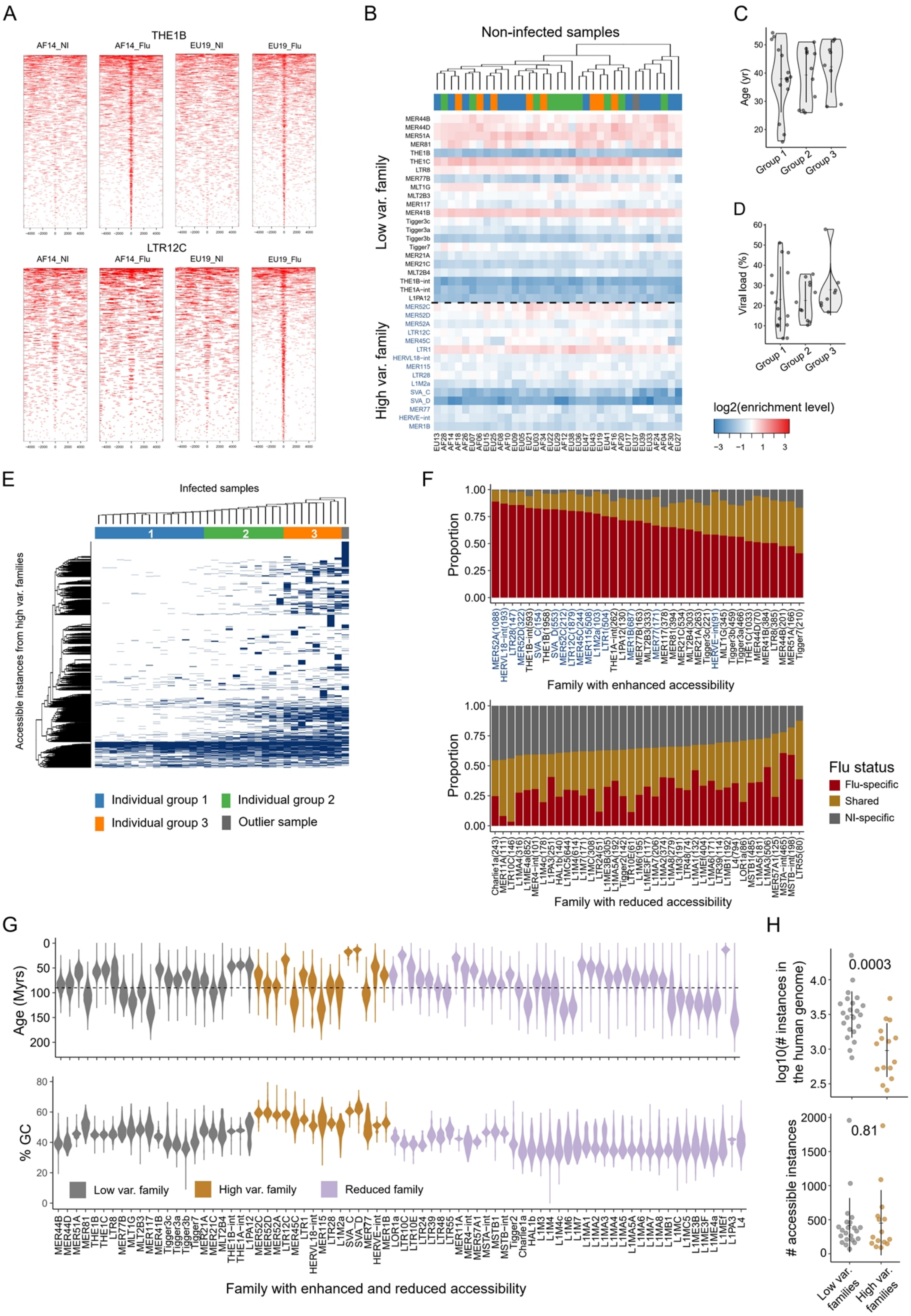
A
THE1B
AF14_NI
AF14_Flu
EU19_NI
EU19_Flu
LTR12C
B
Non-infected samples
Low var. family
High var. family
MER44B
MER44D
MER51A
MER81
THE1B
THE1C
LTR8
MER77B
MLT1G
MLT2B3
MER117
MER41B
Tigger3c
Tigger3a
Tigger3b
Tigger7
MER21A
MER21C
MLT2B4
THE1B-int
THE1A-int
L1PA12
MER52C
MER52D
MER52A
LTR12C
MER45C
LTR1
HERVL18-int
MER115
LTR28
L1M2a
SVA_C
SVA_D
MER77
HERVE-int
MER1B
log2(enrichment level)
C
Age (yr)
Group 1
Group 2
Group 3
D
Viral load (%)
Group 1
Group 2
Group 3
E
Infected samples
Accessible instances from high var. families
Individual group 1
Individual group 2
Individual group 3
Outlier sample
F
Proportion
Family with enhanced accessibility
Family with reduced accessibility
Flu status
Flu-specific
Shared
NI-specific
G
Age (Myrs)
% GC
Low var. family
High var. family
Reduced family
Family with enhanced and reduced accessibility
H
0.0003
log10(# instances in the human genome)
0.81
### accessible instances
Low var. families
High var. families

**Figure S3. Characteristics of TE families display high variability in chromatin accessibility post-infection**

**(A)** Heatmaps of open chromatin regions at THE1B and LTR12C (± 5 kb). Two infected and non-infected samples are shown as examples. LTR12C shows a high variability of accessibility between randomly-selected samples post-infection. ATAC-seq peaks are centered at the summit in TEs. 5 kb upstream and downstream regions are shown. Compared to AF14, EU19 displays a higher enrichment at LTR12C but comparable enrichment at THE1B. **(B)** Heatmap of log2 enrichment levels of 37 enhanced families in 35 non-infected samples. Semi-clustering analysis was performed. Three individual groups observed at infected samples are not clustered together. High var. families are highlighted in blue color. **(C,D)** Violin plots of age of macrophage donors and viral load of the three individual groups. Group 3 individuals have relatively older ages and higher viral load compared with Group 1 individuals. The dot represents each individual and the error bar represents the mean value and standard deviation. **(E)** Heatmap of the chromatin state of accessible instances from high var. families in 35 infected samples. The state of open chromatin is in blue color and closed chromatin in white color. Unsupervised clustering analysis was performed and the three individual groups are clustered together. A fraction of instances shows an enrichment in group 3 compared to group 1 individuals. **(F)** Proportions of fu-specific, shared, and NI-specific instances of each family with enhanced and reduced accessibility. Flu-specific instances represent instances that are accessible in ≥ 1 infected and no non-infected sample; NI-specific instances represent instances that are accessible in ≥ 1 non-infected and no infected sample; Shared instances represent instances that are accessible in ≥ 1 non-infected and ≥ 1 infected samples. High var. families are in blue color. **(G)** Violin plots of the estimated TE evolutionary ages (up) and GC contents (bottom). Dotted lines indicate the evolution time when primates diverged from other mammals (~90 million years ago). No distinct patterns are observed between high var., low var. families, and reduced families. The estimation of ages was described in Methods. High var. families show a higher GC content compared to others. GC contents of ATAC-seq reads per sample are also analyzed (data not shown). Group 3 individuals are observed to have comparable or lower GC contents than other individuals, supporting that the higher accessibilities in high var. families for Group 3 individuals are not derived from sequencing artifacts. **(H)** Number of instances and accessible instances from high var. and low var. families. *P* values computed by two-tailed student's *t*-test are shown above the dot plots.

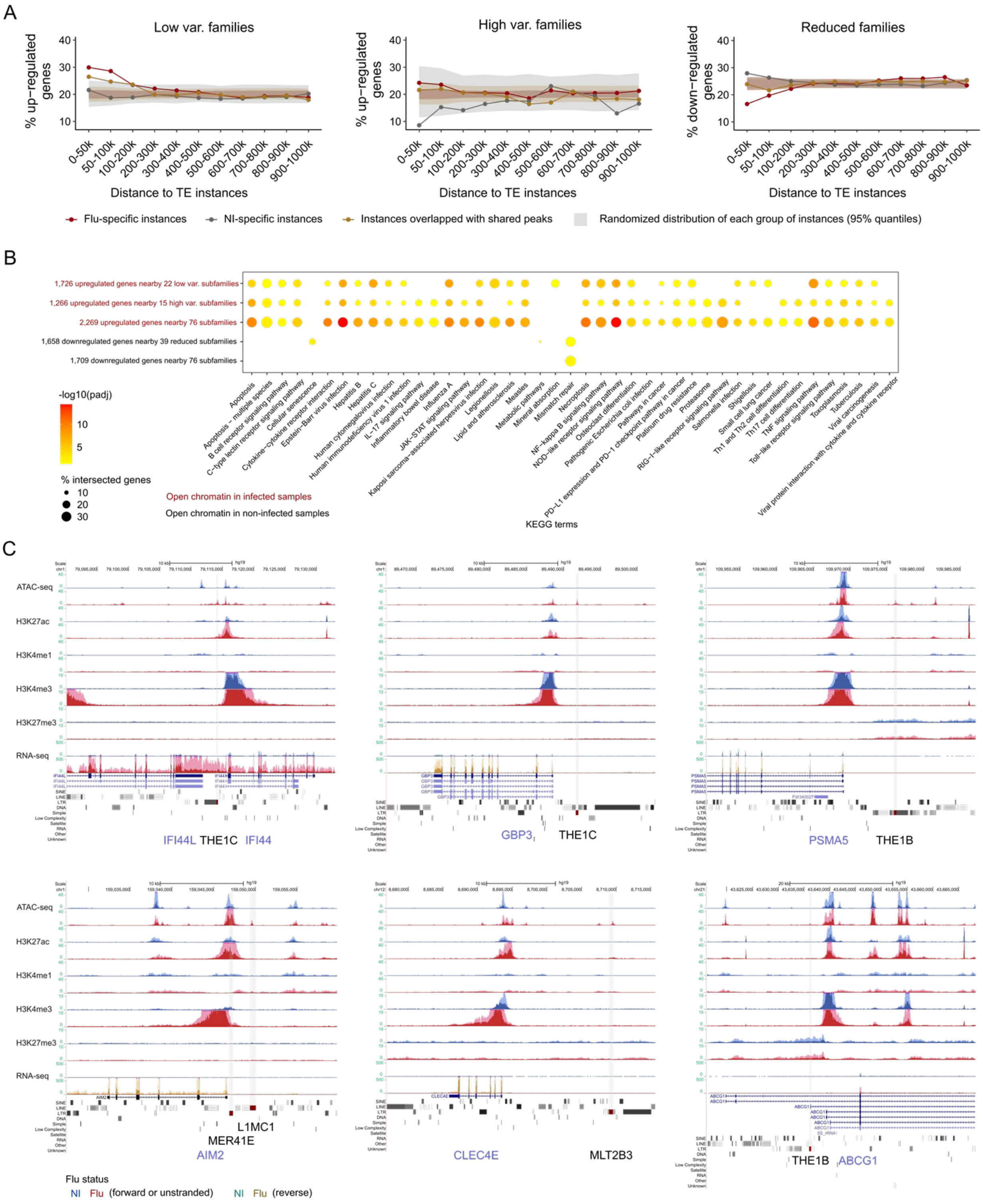
A
Low var. families
High var. families
Reduced families
% up−regulated genes
% down−regulated genes
Distance to TE instances
0-50k
50-100k
100-200k
200-300k
300-400k
400-500k
500-600k
600-700k
700-800k
800-900k
900-1000k
Flu-specific instances
NI-specific instances
Instances overlapped with shared peaks
Randomized distribution of each group of instances (95% quantiles)
B
1,726 upregulated genes nearby 22 low var. subfamilies
1,266 upregulated genes nearby 15 high var. subfamilies
2,269 upregulated genes nearby 76 subfamilies
1,658 downregulated genes nearby 39 reduced subfamilies
1,709 downregulated genes nearby 76 subfamilies
-log10(padj)
% intersected genes
Open chromatin in infected samples
Open chromatin in non-infected samples
KEGG terms
C
ATAC-seq
H3K27ac
H3K4me1
H3K4me3
H3K27me3
RNA-seq
IFI44L THE1C IFI44
GBP3 THE1C
PSMA5 THE1B
L1MC1
MER41E
AIM2
CLEC4E
MLT2B3
THE1B ABCG1
Flu status
NI Flu (forward or unstranded)
NI Flu (reverse)

**Figure S4. TE families with accessibility changes may co-opt in the immune response to IAV infection**

**(A)** Proportions of up/down-regulated genes near flu-specific, NI-specific, and shared instances. Proportions were computed within each of the genomic intervals relative to nearby accessible instances. Upregulated genes were analyzed for high var. and low var. families, and downregulated genes were analyzed for reduced families. Expected distributions were computed as we described in Methods (shaded regions, 95% confidence intervals). The proportions are compared with corresponding expected distributions. Flu-specific instances from low var. and high var. families and NI-specific instances from reduced families display the highest proportions of up/down-regulated genes within 100 kb relative to nearby accessible instances. **(B)** Pathway enrichment analysis of DEGs adjacent (≤ 100 kb) to each category of families. It shows the enrichment of multiple immune-related pathways near families with enhanced accessibility. Some pathways are differentially enriched between high var. and low var. families, including RIG-I-like receptor signaling pathway. **(C)** Example genomic views of instances with enhanced accessibility post-infection. Instances are highlighted as the shaded areas. Six TE immune-related gene pairs are shown, *i.g.*, THE1C-*IFI44*, THE1C-*GBP3*, THE1B-*PSMA5*, MER41-*AIM2*, and MLT2B3-*CLEC4E*, THE1B-*ABCG1*. *AIM2* has been validated to be regulated by a MER41 instance (Chuong et al., 2016); interestingly, it may also be regulated by another TE instance. Other three TE instances reported by Chuong et al. that potentially regulate *APOL1*, *IFI6*, and *SECTM1* did not show chromatin change in macrophages (https://computationalgenomics.ca/tools/epivar). The dark shaded area denotes the distribution of the average RPM values and the light shaded area denotes the standard deviation. Signals of various epigenetic marks are shown in blue color for non-infected samples and red color for infected samples. For RNA-seq, forward and reverse transcripts are shown in blue and green color separately for non-infected samples; while forward and reverse transcripts are shown in red and brown color separately for infected samples.

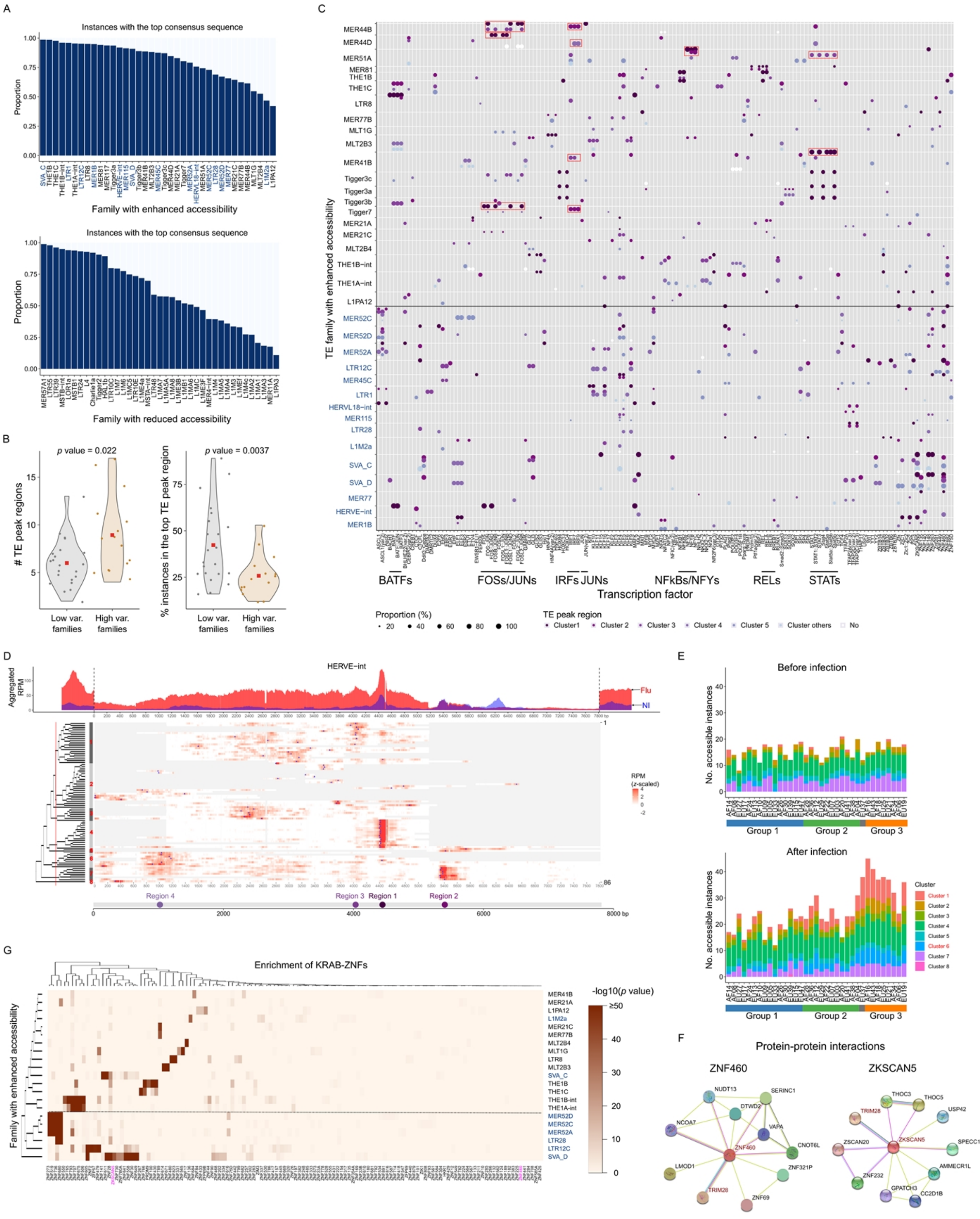

A
Instances with the top consensus sequence
Proportion
Family with enhanced accessibility
Instances with the top consensus sequence
Proportion
Family with reduced accessibility
B
p value = 0.022
### TE peak regions
Low var. families
High var. families
p value = 0.0037
% instances in the top TE peak region
C
TE family with enhanced accessibility
BATFs
FOSs/JUNs
IRFs
JUNs
NFkBs/NFYs
RELs
STATs
Transcription factor
Proportion (%)
TE peak region
D
HERVE-int
Aggregated RPM
Flu
NI
RPM (z-scaled)
Region 4
Region 3
Region 1
Region 2
E
Before infection
No. accessible instances
Group 1
Group 2
Group 3
After infection
Cluster
F
Protein-protein interactions
ZNF460
ZKSCAN5
G
Enrichment of KRAB-ZNFs
Family with enhanced accessibility
-log10(p value)

**Figure S5. TE peak regions and TF binding motifs reveal the association of high variability in chromatin accessibility with KRAB-ZNFs**

**(A)** Proportions of accessible instances with top candidate consensus sequences. Consensus sequence information was achieved from the ".align" files generated by RepeatMasker. It shows a high proportion for families with enhanced accessibility. High var. families are highlighted in blue color. **(B)** Number of TE peak regions detected for high var. and low var. families. Compared to low var. families, high var. families have significantly more TE peaks regions and more instances in the top TE peak region. *P* values were computed by the two-tailed student's *t*-test. **(C)** TF binding motifs enriched in each TE peak region of families with enhanced accessibility. Top five TE peak regions are shown. Instances in other regions and instances not within TE peak regions were analyzed separately. TE peak regions of each family are shown as separate rows. Dot size refers to the proportion of instances in each region containing each motif. Dotted line separates high var. and low var. families and high var. families are also highlighted in blue color. **(D)** Aggregated RPM (up) and RPM (bottom) values on each instance along the HERVE-int consensus sequence. Infected (red) and non-infected (blue) samples are shown separately including the upstream and downstream regions (± 20% of the consensus sequence length). Computed RPMs were *z*-scaled in the heatmap while values below zero are in white color. Deletions relative to the consensus sequence are shown in grey color. Unsupervised clustering analysis was performed with the scaled RPM values to determine the main clusters. Blue triangles indicate the peak centroids referring to the highest RPM values. TE peak regions and positions are shown in the bottom (Same as **Figure 5B**). More details were described in Methods. **(E)** Number of accessible instances from each cluster. Infected and non-infected samples are shown separately. Individuals are ordered based on the clustering obtained in **Figure**

**3C**. Instances from Cluster 1 and 6, most of which contain TE peak region 3 and 4, are more abundant in Group 3 individuals than Group 1 individuals. **(F)** Protein-protein interactions (PPIs) of ZNF460 and ZKSCAN5 TFs and TRIM28, which is involved in the SUMO system. Both ZNF460 and ZKSCAN5 are shown to have experimentally validated (pink lines) PPIs with TRIM28. ZKSCAN5 is also found to interact with THOC5, which is associated with the viral mRNA exportation from the host cell nucleus (gene ontology annotation). **(G)** Enrichment of Zinc finger (ZNF) protein binding sites in high var. and low var. families in 257 HEK293T cell lines (Imbeault et al., 2017). High var. families MER52s, SVAs, LTR12C, and LTR28 are shown to be highly enriched for ZNF binding sites. Color intensity refers to the -log10 $p$ value for enrichment compared to random distribution and one-tailed fisher's exact test was used in the prior study.

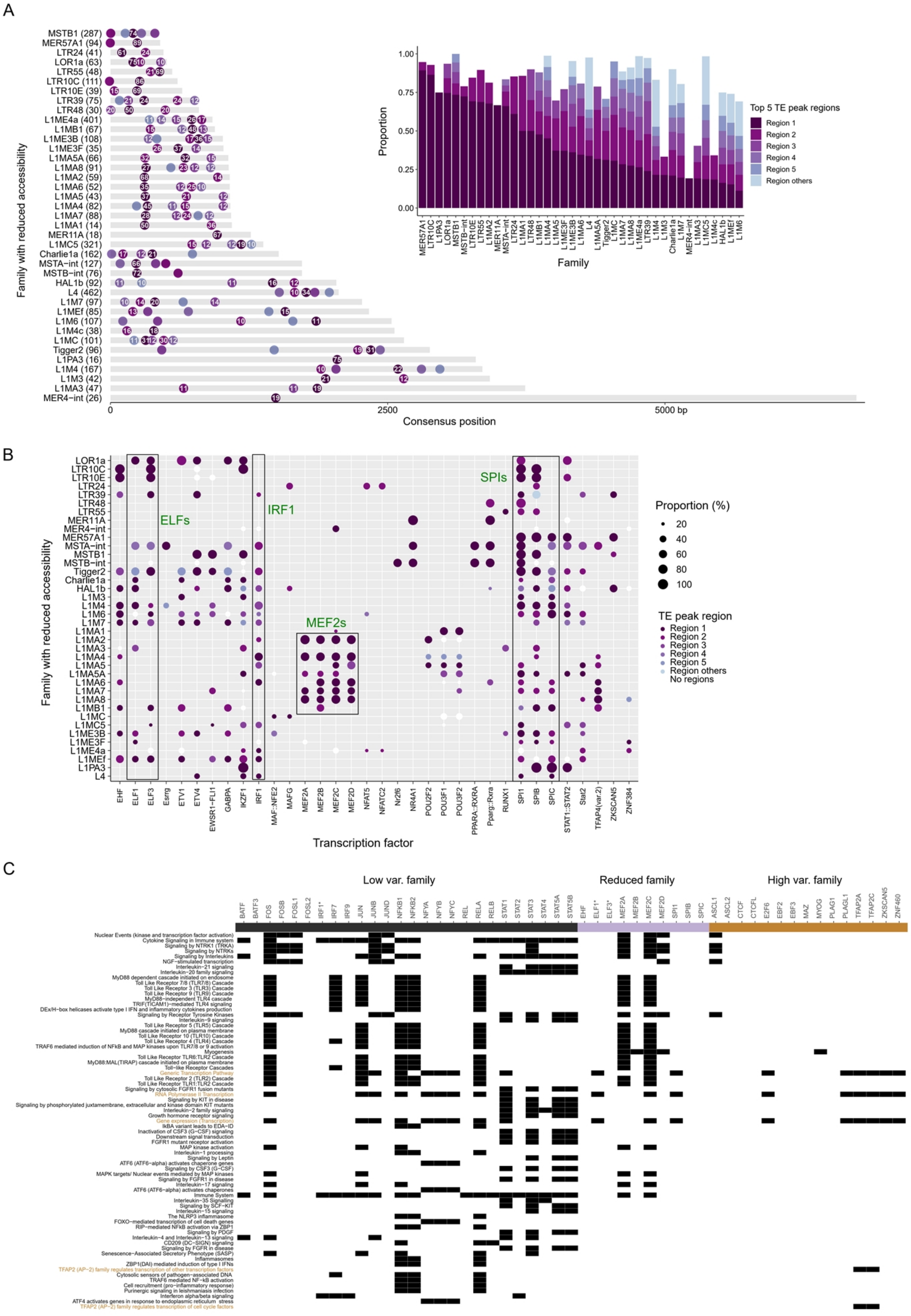
A
Family with reduced accessibility
Consensus position
0
2500
5000 bp
Proportion
Family
Top 5 TE peak regions
Region 1
Region 2
Region 3
Region 4
Region 5
Region others
B
Family with reduced accessibility
Transcription factor
ELFs
IRF1
MEF2s
SPIs
Proportion (%)
20
40
60
80
100
TE peak region
Region 1
Region 2
Region 3
Region 4
Region 5
Region others
No regions
C
Low var. family
Reduced family
High var. family

**Figure S6. TE Peak regions and TF binding motifs enriched in reduced families and different sets of TE families may co-opt in distinct regulatory pathways in response to IAV infection**

**(A)** Distribution and TE peak regions on reduced families. The inset barplot shows the proportion of instances in each TE peak region. The locations and proportions (%) of the top-five TE peaks regions along the consensus sequence are shown. The number in each dot refers to the proportion (above 10%) among accessible instances in each TE peak region. It shows that most instances with reduced accessibility (in Region 1) from L1MA families are consistently located at similar locations of the 3'end of consensus sequence. Y-axis shows the family name and the number of accessible instances mapped to the top consensus sequence. TE peak regions were detected as we described in Methods. **(B)** TF binding motifs enriched in reduced families. Same motifs enriched across TE peak regions were aggregated and the proportions of instances containing each motif are shown. TE peak regions with more accessible instances are shown as representatives. Black boxes highlighted the SPI and MEF2 related motifs (green color) enriched in reduced families. It shows that Region 1s (located at around 300 bp, **Figure S6B**) of L1MA families are consistently enriched for MEF2 related motifs. MEF2 related TFs were previously reported to regulate anti-microbial genes (Clark et al., 2013). **(C)** TFs potentially bound to different categories of families are involved in distinct pathways. Apart from AP-2 related pathways, TFs bound to high var. families are mainly involved in transcription-related pathways. TFs bound to low var. families and reduced families are mainly involved in cytokine signaling and other immune-related pathways. Bars in different colors represent different categories of families they are potentially bound to. * indicates motifs that are enriched in different categories of families. IRF1 motif is enriched in both low var. families and reduced families. ELF1/3 motifs are enriched in both high var. families and reduced families.

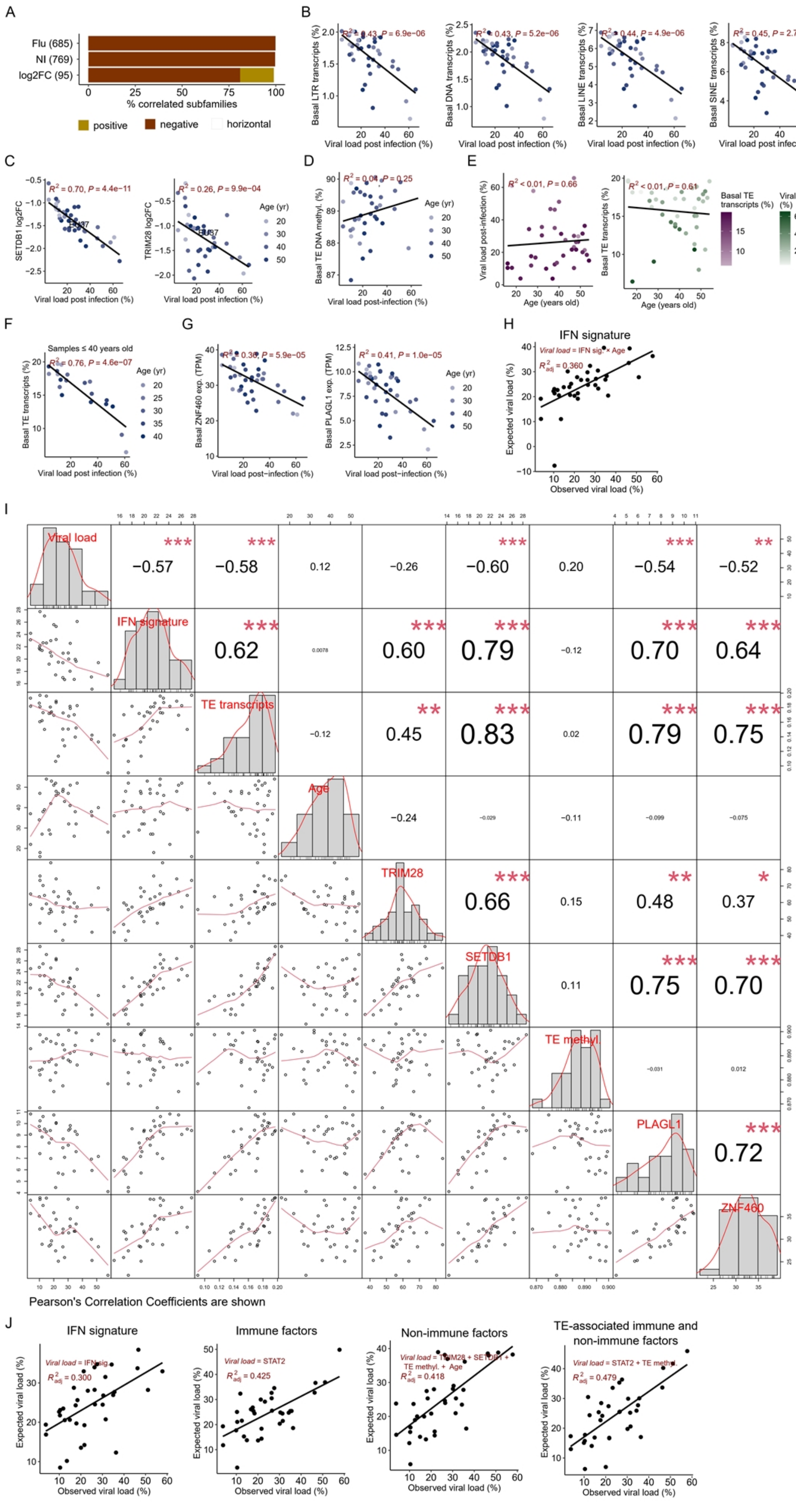

**Figure S7. Correlations between TE-associated host factors and viral load post-infection**

**(A)** Correlation directions among TE families that are correlated ($R^2 \geq 0.3$ and $p$ value $\leq 0.05$) with viral load post-infection (**Figure 6A**). More details were described in Methods. **(B)** Consistent inverse correlations between the basal transcripts of four main TE subclasses and viral load, including LTR, DNA, LINE, and SINE. The basal amount of transcript refers to the proportion of aggregated normalized read counts in each subclass among the global transcripts. Black line represents the regression line. $R^2$ and $p$ values computed by the linear regression model are shown. **(C)** Correlations between the expression fold changes (log2FCs) of both SETDB1 (left) and TRIM28 (right) and viral load. SETDB1 rather than TRIM28 is shown to be inversely correlated with viral load. **(D)** Correlation between the basal DNA methylation level in TEs and viral load. Individuals with low viral load are likely to have lower TE DNA methylation levels. **(E)** Correlations between both viral load post-infection (left) and basal TE transcripts (right) and ages. Samples from younger individuals have a relatively higher amount of basal TE transcripts with a smaller deviation compared to older samples in macrophages. Two samples were identified as outliers having among the lowest amount of basal TE transcripts; strikingly, they also preserved among the highest viral load. **(F)** Correlation between the basal TE transcripts and viral load for individuals 40 years old or younger. It shows an increased correlation compared to the correlation among 39 samples (**Figure 6B**). **(G)** Correlations between the basal expression of two candidate host factors and viral load post-infection. Basal ZNF460 (left) and PLAGL1 (right) expressions are both inversely correlated with viral load post-infection. **(H)** Multivariable regression model developed for the predictive of viral load post-infection using type I interferon (IFN) signature and age only. Details were described in Methods. **(I)** Correlation matrix chart of viral load and variables that are potentially associated with IAV infection and TEs. Histograms and kernel density overlays of each variable are shown. Scatterplot matrix and absolute correlations between each of two variables and viral load are also shown. Red lines represent the distribution. R *chart.Correlation* function was used for the analysis. Pearson's correlation coefficients are shown ($^{*}\ p \leq 0.05$, $^{**}\ p \leq 0.01$, $^{***}\ p \leq 0.001$).

**(J)** Multivariable regression models developed for the predictive of viral load post-infection while age was included as an independent variable. We used the same sets of variables for the models included in **Figure S7H** and **Figure 6H-J**. The adjusted $R^2$ is significantly lower than the model developed by the inclusion of age as an interaction term variable. Details were described in Methods.

#### SUPPLEMENTAL TABLE DESCRIPTIONS

**Table S1.** Samples and sequencing datasets included in this study. Computed viral load, ages, and ancestry information were also included.

**Table S2.** Significantly up/down regulated TE families in response to influenza infection. Details are described in Methods.

**Table S3.** Correlations between TE families and viral load post-infection. The TE basal expression levels, expression levels post-infection, and the expression changes were correlated with viral load separately.

**Table S4.** Number and proportion of ATAC-seq and Chip-seq (H3K27ac, H3K4me1, H3K4me3 and H3K27me3) peaks overlapped with TEs. Total number of paired-end reads and peaks are also included.

**Table S5.** TE families with enhanced or reduced accessibility. Average and standard deviation of the normalized number of peaks-associated instances in infected and non-infected samples were included. Average, standard deviation and coefficient of variation of enrichment levels relative to the background were also included. Average fold enrichment and adjusted *p* values (two-tailed paired student *t*-test, * $p \leq 0.05$, ** $p \leq 0.01$, *** $p \leq 0.001$) between infected and non-infected samples were also included.

**Table S6.** Type I interferon signaling pathways-related genes used to compute the interferon signature (score). The median value of basal expression levels (TPMs) of 39 genes was used as the interferon signature.
